# Domain-adversarial learning predicts clinically actionable drug combination synergy in leukemia patients using bulk transcriptomics data

**DOI:** 10.64898/2026.05.18.725869

**Authors:** Jie Zhu, Weikaixin Kong, Thi Huong Lan Do, Sandra Kummer, Jarno Kivioja, Rafael Romero-Becerra, Juho Rousu, Mitro Miihkinen, Jeffrey W Tyner, Thorsten Zenz, Tero Aittokallio

## Abstract

Deep learning has gained popularity in drug combination synergy prediction; however, DL models require large training datasets from cell line pharmacogenomic screens that poorly capture the heterogeneity in transcriptomic features and phenotypic responses seen in patients. To that end, we developed a domain-adversarial neural network (DANN) for personalized drug synergy prediction that accounts for systematic differences between cell line and patient domains. In applications to AML and CLL patient samples, we demonstrate how DANN boosts prediction accuracy under realistic data constraints. The model predictions demonstrated elevated synergy among clinically used combinations, such as venetoclax-based regimens, supporting its ability to identify both pharmaceutically and clinically meaningful combinations. DANN estimates prediction uncertainty and prioritizes high-confidence combination predictions to aid clinical translation. Together, DANN provides a systematic approach to improving accuracy and reliability in cross-domain drug synergy prediction, advancing the development of methods that are aligned with the translational requirements of precision hematology.

## Introduction

Combination therapies play a central role in contemporary cancer treatment^1,2^. Patient-matched drug combinations improve therapeutic efficacy, counteract resistance mechanisms, and reduce dose-related toxicities compared to single-agent regimens^3,4^. In hematological malignancies, such as acute myeloid leukemia (AML) and chronic lymphocytic leukemia (CLL), patient-specific tailoring of synergistic drug combinations is particularly critical, as patient responses are highly heterogeneous and often poorly captured by monotherapy response biomarkers^5^. Machine learning offers a systematic approach to navigating the massive combinatorial search space with the aim to predict most synergistic combinations for each patient. Although high-throughput drug combination screens in cancer cell lines have generated large data resources for development of synergy prediction models^6–13^, translating these models from *in vitro* preclinical systems to patient-derived samples and clinical outcomes remains a major unresolved challenge.

A central translational barrier lies in the systematic domain differences between cancer cell lines and primary patient samples^14,15^. Immortalized cell line cultures are relatively homogeneous *in vitro* systems, whereas fresh patient samples are shaped by microenvironmental influences, clonal diversity, and cellular composition^16–19^. Consequently, discrepancies arise not only in genetic and molecular distributions, but also in drug response and combination synergy phenotype distributions^20,21^. ML models trained exclusively on cell line data may therefore learn response patterns that are biased toward the source domain and fail to generalize to patient context, limiting their translational utility. Drug sensitivities tested directly in patient cells (*ex vivo* drug testing) can be used for personalized combination selection, but these models require additional molecular data, such as single-cell transcriptomics^22,23^, that are often not routinely profiled for each patient in clinical settings.

Recent efforts have sought to address this cross-domain gap by using patient data as part of model training or by applying transfer learning strategies^21,24–27^. These approaches implicitly assume that drug combination synergy manifests in comparable statistical properties across the domains and focus primarily on aligning molecular feature representations between cell lines and patient samples. However, drug responses in patient-derived cells are markedly more heterogeneous and context-dependent than those observed in cell lines^28^, resulting in potential distribution shifts in the prediction target that are not explicitly modeled by the existing methods. Such domain-dependent differences in the outcome distribution may systematically bias predictions, even when the molecular feature alignment is partially achieved. Addressing domain shifts both at the level of input representation and prediction target is therefore essential for robust translational modeling.

These challenges are particularly important when predicting drug combination responses in patients with relatively rare leukemia types such as AML and CLL. High patient-to-patient heterogeneity in response patterns and a limited availability of preclinical drug combination data and patient clinical outcomes reduces accuracy, robustness and clinical applicability of the models that require large-scale multi-domain training datasets. Furthermore, the absence of uncertainty estimates poses translational challenges to clinical interpretability; models that achieve high average accuracy across patients may still fail on atypical patients or extreme responses, while providing no indication of when predictions should be treated with caution. These limitations underscore the need for practical prediction approaches that are robust to domain shifts and provide accurate predictions with confidence estimates under realistic data constraints in personalized hematology settings.

Here, we develop a domain-adversarial neural network (DANN) framework for drug combination synergy prediction that leverages limited blood cancer cell line data for pre-training on multi-modal molecular features, including drug-induced gene expression signatures, chemical structure fingerprints, and pathway-level activity scores. The model is then adapted to patient combination response data using adversarial learning, in which a shared feature extractor is optimized to support accurate synergy prediction, while simultaneously discouraging domain-specific representations. This learning strategy promotes domain-invariant and task-relevant embeddings, mitigating biases arising from shifts in both transcriptomic feature and synergy distributions. Applied to patients with AML and CLL, we show how the DANN framework generalizes across biological and clinical contexts across distinct leukemia types, requiring only minimal numbers of both cell lines and patient samples tested with drug combinations for accurate predictions that are well-aligned with established therapeutic efficacy in AML and CLL clinical management.

## Results

### A domain-adversarial framework transfers synergy prediction from cell lines to patients

We developed a DANN approach to predicting drug combination synergy for each patient sample separately, across a wide range of potential drug combinations, based on multi-modal molecular features. The framework uses a two-stage training strategy comprising cell line pre-training followed by patient-specific domain adaptation (**Fig. 1A**).

**Figure 1.**
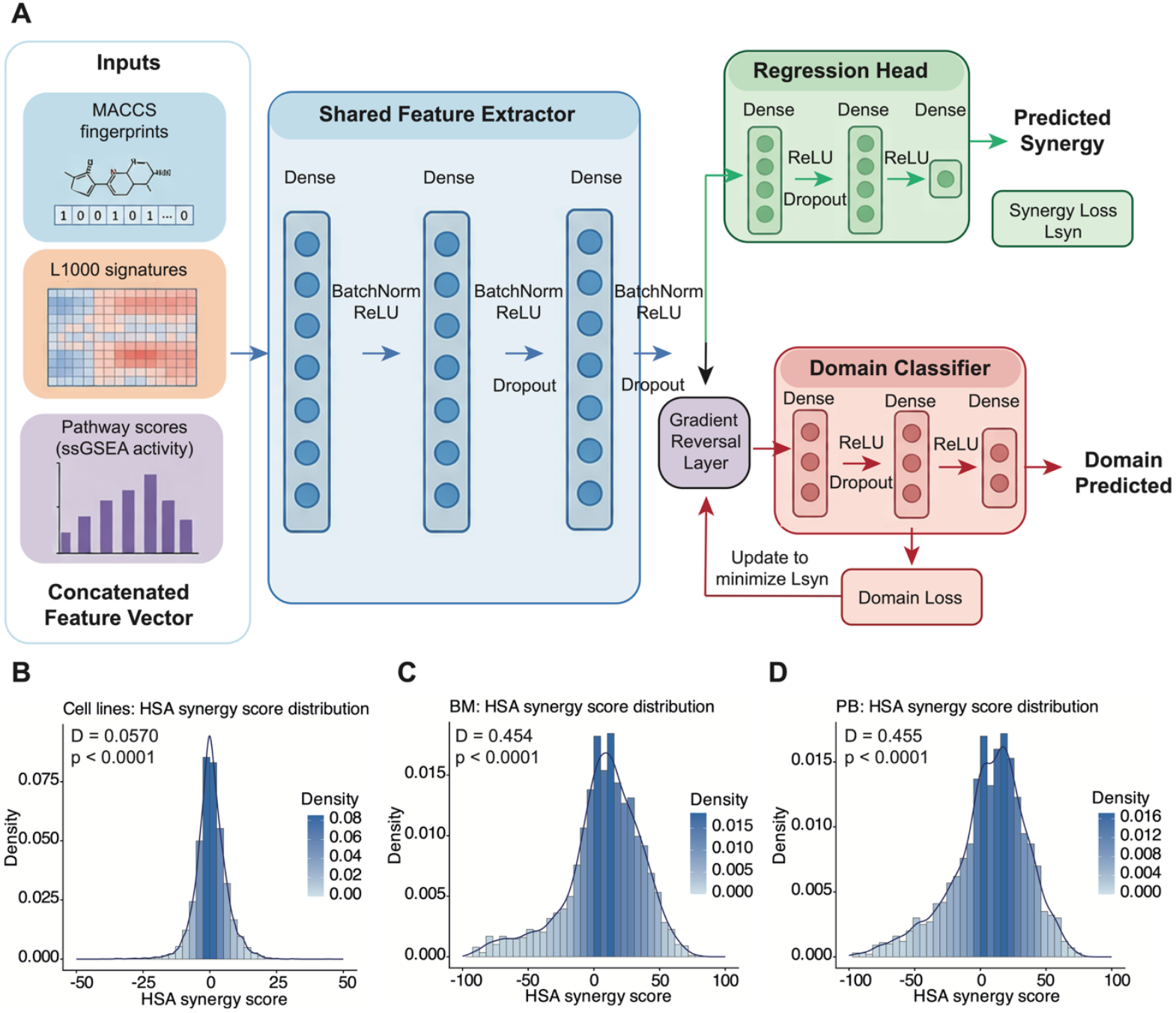
Domain-adversarial framework for drug synergy prediction accounts for domain shift between cell lines and patient samples. (**A**) Architecture of the domain-adversarial neural network (DANN) model, which consists of a shared feature extractor that learns latent representations from multi-modal drug features, including chemical structure, transcriptional perturbation, and pathway activity. The extracted features are fed into two branches: a regression head for drug synergy prediction and a domain classifier for distinguishing between cell line and patient samples. A gradient reversal layer is inserted before the domain classifier to enable adversarial training, encouraging the learned features to be domain-invariant while preserving predictive performance. (**B**) Distribution of HSA synergy scores of 2,150 combinations tested in nine blood cancer cell lines, showing a relatively narrow combination response range centered around additive effects, significantly deviating from a normal distribution (Kolmogorov–Smirnov test, *p* < 0.0001). (**C**) Distribution of HSA synergy scores of 55 combinations tested in 72 AML patients’ bone marrow (BM) samples used for model fine-tuning, exhibiting a wider combination response range and a shift toward positive synergy values, different from that of cell line synergies (*p* < 0.0001). (**D**) Distribution of HSA synergy scores of 55 combinations tested in 74 AML peripheral blood (PB) samples held-out for testing, showing a similarly widened and positively skewed distribution than BM samples, and difference from the cell lines (*p* < 0.0001), while no significant difference between AML BM and PB sample distributions (*p* = 0.53).

In the pre-training stage, the model was trained exclusively on *in vitro* data from nine blood cancer cell lines to learn a generalizable relationship between drug-induced molecular perturbations and drug combination synergy outcomes (**Suppl. Table 6**). The architecture consists of a shared feature extractor that integrates three complementary inputs: (i) L1000 differential gene expression signatures induced by single agents (1,104 compounds), (ii) MACCS fingerprints encoding the chemical structures of each compound pair, and (iii) ssGSEA pathway scores computed separately from baseline gene expression data of cell lines and patient samples before drug treatments. These features were concatenated and passed through a neural network to predict drug combination synergy (continuous HSA, Bliss and ZIP synergy scores). Importantly, no patient-derived samples were used during this pre-training phase.

During the domain adaptation, a patient-specific regression model was trained jointly with a domain classifier that discriminates between cell line and patient samples. The shared feature extractor was optimized adversarially to confuse the domain classifier, while simultaneously supporting accurate synergy prediction through the patient regression head. Domain adaptation and fine-tuning were performed separately for AML and CLL patient cohorts to account for disease-specific biological contexts using bulk RNA-seq transcriptomic data that are routinely available from each patient sample. The DANN framework was designed to mitigate the distributional mismatch between cell line and patient samples by learning domain-invariant, yet task-relevant representations, thereby improving cross-domain generalization from cell lines to patient-derived contexts.

In AML patients, the model was fine-tuned using *ex vivo* drug combination response data from bone marrow (BM) samples (*n* = 72), where subsets of varying sizes were used for domain adaptation, and peripheral blood (PB) samples from independent patients (*n* = 74) were held-out exclusively for model testing. This setup provides a stringent evaluation of cross-compartment generalization between BM and PB specimens. In CLL patients, the model was similarly fine-tuned using subsets of PB samples (*n* = 62) with varying training sample sizes, while the remaining samples were reserved for model evaluation. As all CLL data are patient-derived (no CLL cell lines in the pre-training datasets), this setting primarily assesses generalization to unseen patients and disease context, while the principal domain shift arises from the transition between cell line pre-training and patient-derived fine-tuning.

### Significant distributional shifts in drug combination effects between cell lines and patient samples

Compared with the cell line combination responses, the AML patient-derived samples exhibited significantly different drug synergy distributions both for BM and PB specimens (*p* < 0.0001, Kolmogorov–Smirnov test), while the distributions of BM and PB samples were similar (*p* = 0.53; **Fig. 1B–D**). Consistently, the proportion of synergistic combinations (HSA > 0) was significantly higher in patient-derived samples than in cell lines (*p* < 0.0001, Fisher’s exact test; odds ratio (OR) = 0.55 for BM and 0.59 for PB; **Fig. 1C-D**). Such differences in the scale and shape of the target distribution imply that models trained only on cell line data may become biased toward the source-domain response profile and therefore misestimate synergy in patient samples. Similar domain-dependent shifts were observed for the alternative synergy metrics (Bliss and ZIP; **Suppl. Fig. 1A-F**).

Notably, patient samples displayed increased dispersion and heavier synergistic and antagonistic tails compared with cell line distributions (**Fig. 1C-D**), indicating a greater heterogeneity in drug combination responses across individual patients. Subsampling of the patient samples confirmed that the differences were not due to differences in the sample sizes (**Suppl. Fig. 1G-I**). A degree of enrichment of synergistic combinations is expected in patient samples, since scarce patient primary cells are often tested with combinations that are expected to show synergy; in contrast, *in vitro* cell line screens can be performed in a more random set of drug combinations. This wider synergy range in patient samples further underscores the presence of a domain shift that affects not only the molecular feature distributions but also the scale and variability of the prediction target itself.

### Domain-adversarial learning improves cross-compartment generalization in AML patient samples

We next evaluated the predictive performance of the DANN model in the AML setting, focusing on its ability to generalize from BM samples used for fine-tuning to PB samples held-out for testing; we compared the performance of DANN with four baseline models trained directly on the patient samples (XGBoost, random forest, k-nearest neighbors and linear regression), across varying sizes of fine-tuning patient samples (**Fig. 2**).

**Figure 2.**
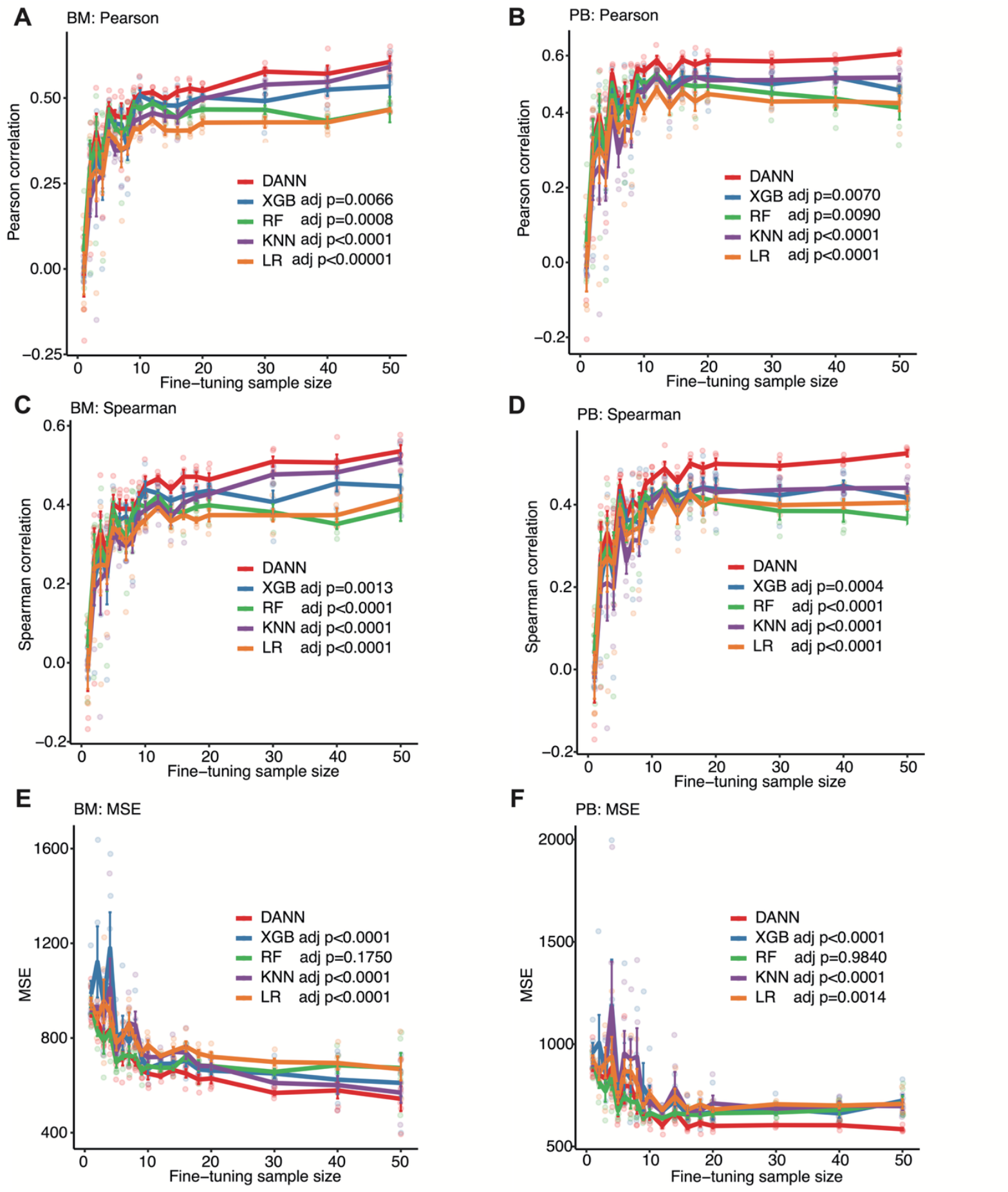
Predictive performance of DANN and baseline models for AML patient-derived *ex vivo* responses. Performance of DANN at predicting HSA synergy compared with the baseline models (XGBoost, random forest, k-nearest neighbors, and linear regression) as a function of the number of fine-tuning samples. Performance was evaluated separately for BM and PB samples using Pearson correlation, Spearman correlation, and MSE. (**A, C, E**) Model performance on the remaining AML BM samples not used in model fine tuning. (**B, D, F**) Model generalization performance on held-out AML PB samples. The lines represent average performance across five repeated sampling runs, with error bars indicating standard error. Significance between the models was assessed via a robust linear mixed-effects model (LMM) to account for inter-iteration variability. Pairwise contrasts between DANN and other models were adjusted for multiple testing using Tukey’s method (**Suppl. Tables 1-2**).

As expected, all models showed rapid performance gains in Pearson correlation on the BM samples as training samples increased (**Fig. 2A**). While performance was unstable with fewer than five BM samples, coefficients gradually plateaued between 0.4 and 0.6. Notably, DANN consistently achieved higher Pearson correlations than baselines across all sample sizes (**Suppl. Table 1**). These trends were mirrored in the generalization to held-out PB samples (**Fig. 2B, Suppl. Table 2**); although all models exhibited limited predictive power in the low-data regimen, DANN maintained a clear advantage, particularly as the number of training samples approached ten. Similar trends were observed for Spearman correlation in the BM and PB domains (**Fig. 2C-D**). We further evaluated models’ performance through mean squared error (MSE). In the BM fine-tuning domain, MSE converged to dataset-specific minima as training reached 30–50 samples (**Fig. 2E**). Across both the BM test samples and the held-out PB samples, DANN consistently achieved the lowest MSE among all tested models (**Fig. 2F**). This advantage was particularly pronounced in the PB samples, where the gap between DANN and the baselines widened as the number of training samples increased.

The benefits of domain-adversarial learning in AML were not limited to HSA-based synergy. When evaluated using Bliss or ZIP synergy scores, DANN again showed improved robustness compared with the baseline models (**Suppl. Figs. 2,3**). To further dissect the contributions, an ablation analysis compared the full DANN model with a patient-only neural network (**Suppl. Fig. 4, Suppl. Table 3**). In the BM fine-tuning domain, the DANN model showed a trend toward higher Pearson correlation both in the BM setting (*p* = 0.056) and the held-out PB samples (*p* = 0.056) (**Suppl. Fig. 4A**-**B**). Although the Spearman ranked correlation showed positive trends that did not reach the traditional significance threshold (**Suppl. Fig. 4C-D, Suppl. Table 3**), the model’s impact on the absolute predictive error was evident in significant reduction in MSE across both BM and PB (*p* < 0.001) samples, compared to the patient-only model (**Suppl. Fig. 4E**-**F**). These results indicate that while both architectures capture ranking and linear trends with a comparable efficacy, DANN significantly improves the absolute accuracy of drug combination synergy predictions.

Overall, these results demonstrated how DANN provides systematic improvements in cross-compartment generalization from cell lines to AML patient samples, particularly under limited training data setting that is frequent in patient-based studies, while preserving competitive performance on the training domain as well.

### DANN generalizes across leukemia types and improves prediction in CLL patient samples

We next evaluated the DANN predictive power on an independent CLL drug combination dataset to assess its generalizability across two distinct leukemia types. This patient dataset consists of *ex vivo* drug synergy measurements of 90 unique combinations in primary CLL patient PB cells (*n* = 62). To evaluate the model performance under varying data availability settings, the pre-trained model was fine-tuned using varying subsets of CLL samples with increasing sample sizes, while the remaining samples were reserved for model evaluation. Since all CLL data are patient-derived, this analysis primarily tests generalization to unseen patients within the same cohort, while the principal domain shift arises from the transition between cell line–based pre-training and patient-derived fine-tuning.

Compared to the cell line responses, CLL samples exhibited a significantly altered drug combination synergy distribution (*p* < 0.0001, Kolmogorov–Smirnov test; **Fig. 3A**). Notably, a pronounced distributional shift was observed also between CLL and AML patient PB samples (*p* < 0.0001), which is likely due to specific experimental design that used relatively low drug doses in the CLL samples, resulting in a decreased synergy scores and a median cell toxicity of approximately 10–20% (**Suppl. Fig. 5**). Across all patient-based fine tuning sample sizes, DANN consistently achieved higher predictive performance than the baseline methods (**Fig. 3B–D**). In the low-sample regimen, all the models showed a relatively limited and unstable performance, indicating a more challenging combination prediction setting, due to the lower absolute values of combination synergy in CLL samples.

**Figure 3.**
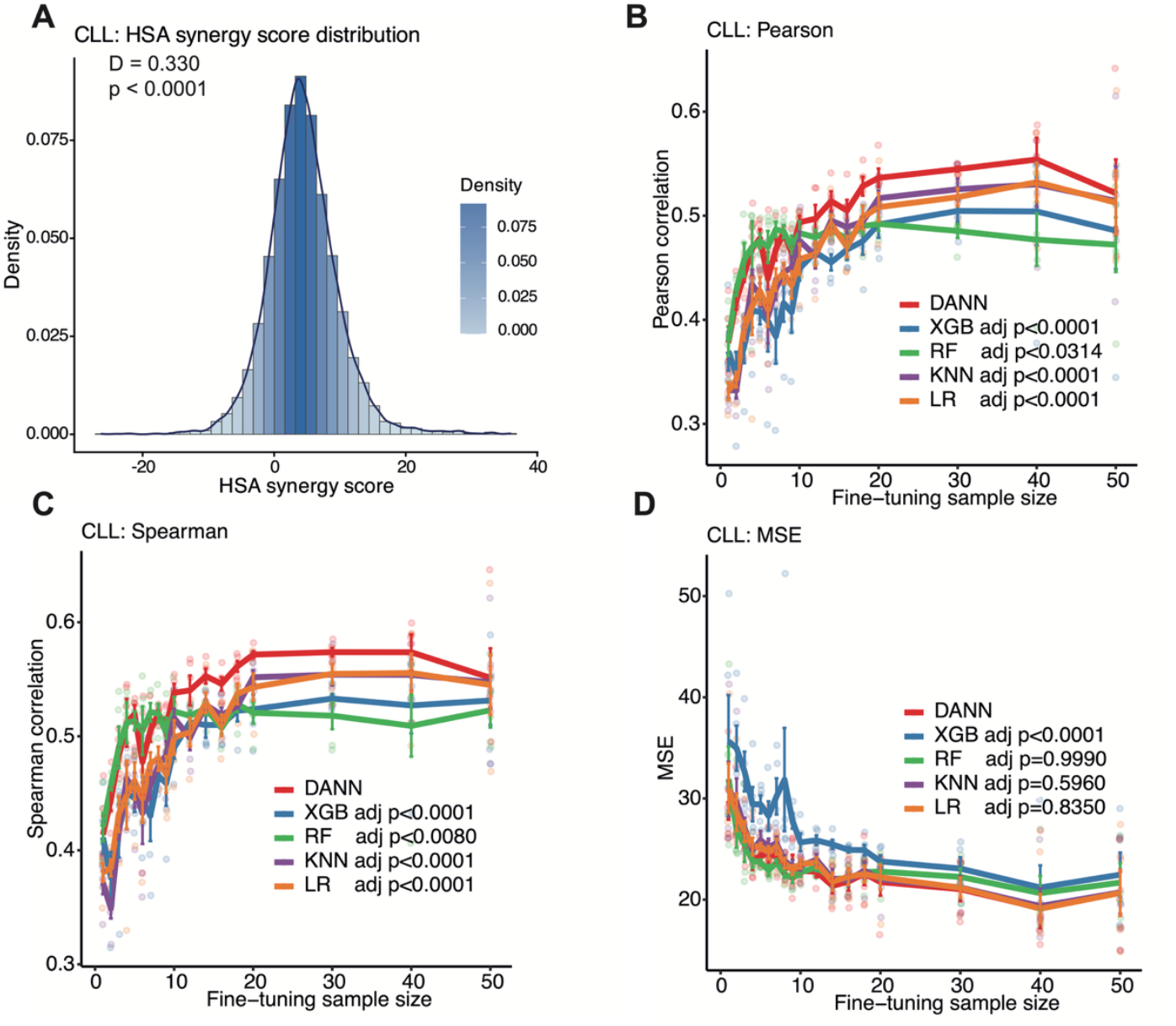
Drug synergy prediction performance in CLL patient PB samples. (**A**) Distribution of HSA synergy scores of 90 combinations in 62 primary CLL patient samples, showing a moderately narrow distribution with a slight shift toward positive synergy values, compared to the cell line synergy distribution (*p* < 0.0001, Kolmogorov–Smirnov test). (**B–D**) Predictive performance of DANN and the baseline models as a function of the number of CLL fine-tuning training samples, evaluated by Pearson correlation, Spearman correlation, and MSE. Model performance was assessed on the remaining CLL samples not included in the training set, with lines showing average performance and error bars standard error across five repeated runs. Significance between the models was assessed via a robust LMM to account for inter-iteration variability. Pairwise contrasts between DANN and other models were adjusted for multiple testing using Tukey’s method (**Suppl. Table 4**).

As the number of CLL training samples increased, performance improved gradually with all the methods. With approximately 16–20 training samples, DANN reached Pearson correlations in the range of 0.50, exceeding the best-performing baseline models (*p* < 0.05; **Suppl. Table 4**), which plateaued at lower correlation levels (**Fig. 3B**). Spearman correlations followed a similar trend, indicating improved ranking of drug combinations by the predicted synergy (**Fig. 3C**). At higher sample sizes (*n* = 20–40), the performance of models stabilized, with DANN achieving moderate correlations and maintaining a consistent advantage over the baseline approaches. At the largest sample size (*n* = 50), the average performance of all models dropped, likely due to increased variability arising from the reduced size of the held-out test set and the inclusion of more heterogeneous patient samples (**Fig. 3D**).

Overall, these results demonstrate that domain-adversarial learning provides robust and reproducible gains also in CLL patient drug synergy prediction, particularly in data-limited settings, and that the benefits observed in AML generalize to a different leukemia type. By leveraging cell line data during pre-training, even without any CLL cell lines present, DANN compensates for the limited patient data and captures pharmacogenomic relationships transferable to primary CLL samples.

### Error-guided selective prediction prioritizes high-confidence drug synergy predictions

While improving model’s predictive accuracy is important, translational applications further require the ability to identify high confident predictions for a given patient and combination. To address this, we developed a *post-hoc* error prediction model that estimates the expected prediction error of DANN and enables selective prediction, whereby only high-confidence predictions are retained for downstream use (**Fig. 4A**).

**Figure 4.**
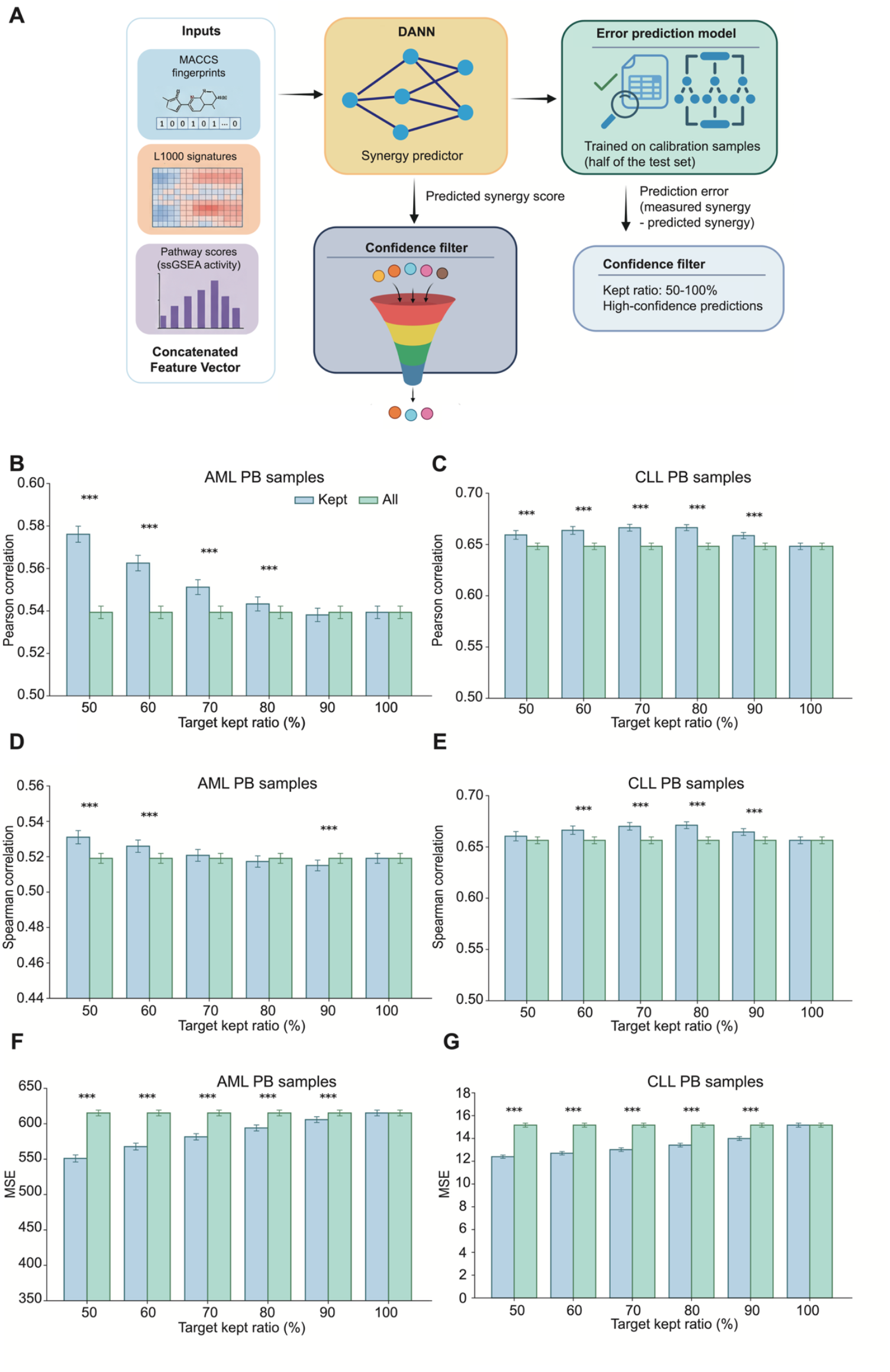
Error-guided selective prediction enhances accuracy in AML and CLL samples. (**A**) Schematic overview of the error-guided selective prediction framework. Drug-induced gene expression signatures, chemical structure features, and pathway activity scores are first used in the DANN model to predict drug combination synergy. A *post-hoc* random forest error prediction model, trained on calibration samples, estimates the expected prediction error for each test sample. Predictions are then ranked by the predicted error and filtered according to a target kept ratio, retaining only high-confidence predictions. (**B, D, F**) Selective prediction performance in the AML PB test cohort (*n* = 37 samples) for the retained (kept) predictions compared with all predictions across different target-kept ratios. Results for the *n* = 11 AML BM samples are shown in Suppl. Fig. 6. (**C, E, G**) Selective prediction performance in the CLL PB cohort (*n* = 6 samples). In all panels, bars represent mean values across repeated sampling runs, with the error bars indicating standard error across the runs. Statistical significance between kept and raw predictions was assessed using the Wilcoxon signed-rank test and the *p*-values were adjusted for multiple comparisons using the Benjamini–Hochberg false discovery rate (FDR) correction; *** adjusted p < 0.001.

Using the AML data cohort as a disease application, we trained a separate random forest regressor to predict the log-transformed absolute prediction error of the DANN model. The error model was trained on half of the test patient cohort using 3-fold cross-validation, separately for the two sample types (*n* = 11 for the BM, *n* = 37 for the PB), and evaluated on the remaining held-out samples (*n* = 11 for BM, *n* = 37 for the PB), ensuring that error estimates generalize across the patients. The input to the error prediction model consisted of the same z-scaled feature representation used by DANN, augmented with the DANN predictions and simple transformations thereof, allowing the model to capture systematic uncertainty patterns associated with either extreme or unstable predictions. Predicted errors were then used to rank test set predictions, and the performance was evaluated after retaining only a specified fraction of predictions with the lowest predicted error (target-kept ratio).

Prioritization of higher-confidence predictions consistently improved performance in the AML PB test set (**Fig. 4B,D,F**). At kept ratios between 50–80%, the retained subset exhibited significantly higher Pearson and Spearman correlations, compared to using all predictions, together with significantly reduced absolute MSE levels. These improvements were statistically significant across most kept ratio thresholds (adjusted *p* < 0.001, Wilcoxon signed-rank test), indicating that the error model effectively identifies predictions that would otherwise contribute disproportionately to the overall error. As the kept ratio approached 100%, performance converged to the original, unfiltered baseline, as expected. Similar trends were observed in AML BM samples (**Suppl. Fig. 6**), supporting the robustness of the approach across sample types.

We next applied the same error prediction and selective evaluation framework to the CLL cohort to assess its robustness across diseases. The error model was trained and evaluated using the same procedure as in the AML case, without modification of model architecture or selection strategy. Prioritization of predictions with lower predicted error led to systematic increases in Pearson and Spearman correlations and lower absolute MSE levels, compared with using all predictions (**Fig. 4C,E,G**). Performance gains were most pronounced at intermediate kept ratios (approximately 60–90%), where improvements were statistically significant across repeated splits. At full coverage (100% kept), the selective and raw performance converged, confirming that gains arise specifically from error-based filtering, rather than from rescaling or re-estimation.

Notably, the magnitude of improvement in CLL samples was comparable to, and in some cases even larger than that observed in AML, despite differences in disease biology and data distributions. This indicates that the learned error patterns are not disease-specific, but instead capture more general uncertainty signals associated with cross-domain synergy prediction. Together, these results demonstrate that the error prediction module provides a practical mechanism to prioritize most reliable predictions for individual patients, enabling a trade-off between prediction coverage and accuracy that is particularly relevant for clinical decision-making.

### Model predictions recapitulate elevated synergy of clinically approved drug combinations

To assess whether the model captures clinically relevant drug combination effects, we compared the predicted synergy scores of clinically successful combinations with those of randomly sampled drug pairs tested in the patient cohorts. We identified clinically approved combinations tested in AML and CLL as referenced in FDA approval summaries and regulatory documentation from finished clinical trials demonstrating clinical benefit (**Suppl. Table 5**). Since azacitidine, a hypomethylating agent used to treat patients with AML, is not represented in the L1000 dataset, we identified structurally similar compounds that could serve as its surrogates. Using the MACCS fingerprint–based Tanimoto similarity, we identified three compounds with high structural similarity to azacitidine (Tanimoto > 0.9): fazarabine, cytarabine, and decitabine.

Among these, venetoclax + decitabine exhibited high predicted synergy scores in the BeatAML cohort significantly shifted toward higher values compared with random combinations (**Fig. 5A-B, Suppl. Fig. 7A-D**). This combination, in which the BCL-2 inhibitor venetoclax is combined with the hypomethylating agent decitabine, has demonstrated efficacy and tolerability in elderly patients with AML^29^, with additional benefits also reported in early-phase trials for high-risk myelodysplastic syndrome^30^. Similarly, venetoclax + cytarabine, a regimen used for AML patients ineligible for intensive chemotherapy^31^, demonstrated a significant increase in predicted synergy (*p* < 0.0001; **Fig. 5C-D, Suppl. Fig. 8A-D**). While the synergy metrics HSA, ZIP and Bliss varied in their absolute values, the consistent enrichment of higher synergies highlights the model’s capacity to identify therapeutically successful combinations. In the venetoclax + fazarabine combination, the results were mixed; while no significant differences were observed compared to random pairings using HSA and ZIP, a significant increase in synergy was predicted by the Bliss model (**Suppl. Fig. 9A-F**).

**Figure 5.**
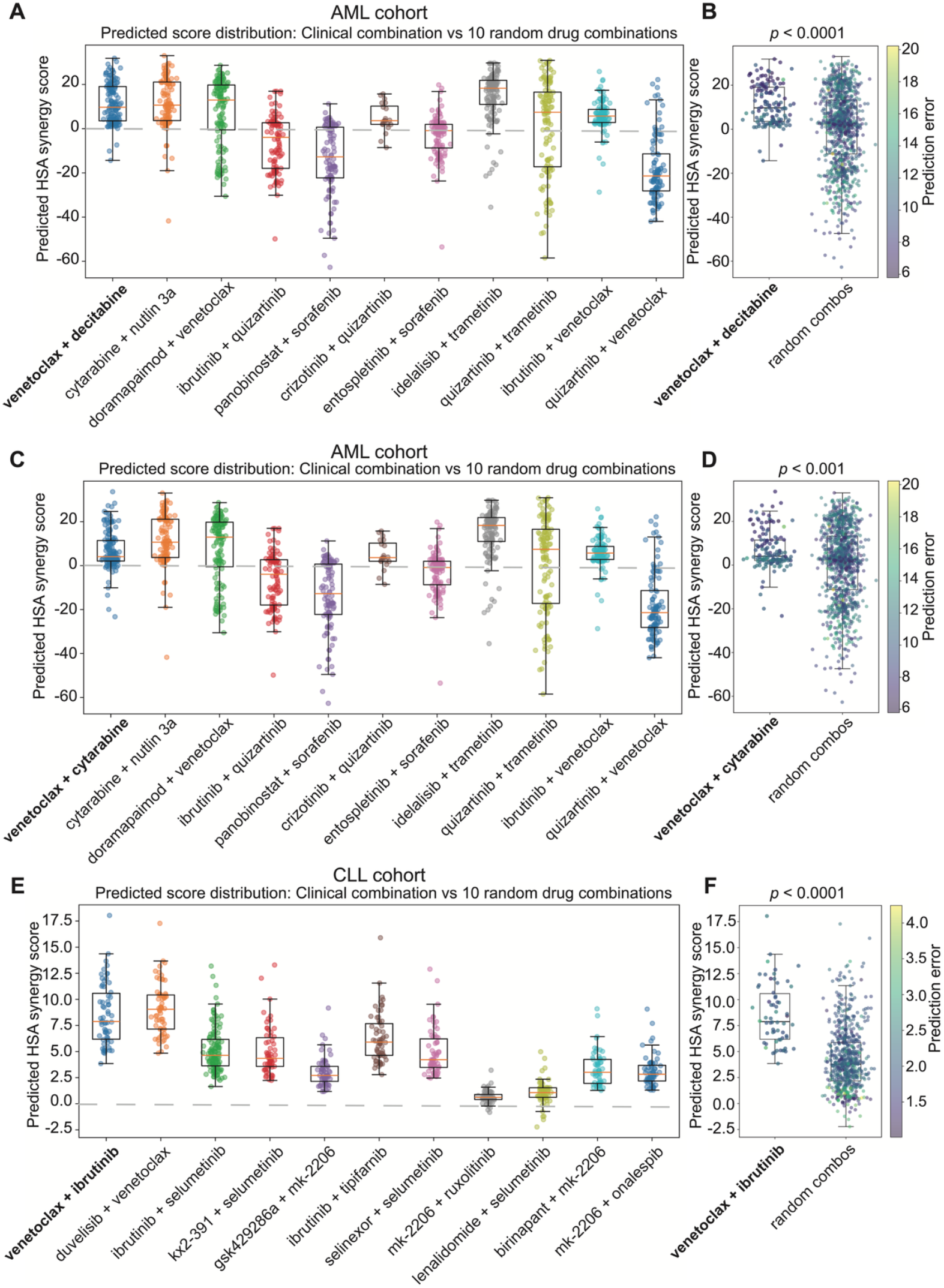
Model predictions show elevated synergy in clinically successful drug combinations. Distribution of predicted synergy scores for the clinically approved combinations (bold): (**A**) venetoclax + decitabine in the AML cohort, (**C**) venetoclax + cytarabine in the AML cohort, (**E**) venetoclax + ibrutinib in the CLL cohort, compared with ten randomly sampled drug combinations among those tested in AML samples and CLL samples, respectively. The clinical combinations exhibited a clear shift toward higher predicted synergy values relative to the random combinations. (**B, D, F**) Statistical comparison of predicted synergy scores between the clinical combinations and random drug combinations in AML samples (*n* = 146, including both BM and PB samples) and CLL samples (*n* = 62). Each point corresponds to an individual patient prediction, colored by prediction confidence, and box plots show the median (horizontal line) and the interquartile range (boxes). Statistical significance was assessed using a two-sided Mann–Whitney U test.

We next evaluated the clinically approved AML combinations of venetoclax + decitabine and venetoclax + cytarabine in the CLL cohort. Consistent with the AML results, venetoclax combined with either decitabine or cytarabine showed higher predicted synergy scores, when compared with random drug combinations (**Suppl. Fig. 10A-D**). We further tested the clinically used combination of venetoclax and ibrutinib, which has a strong clinical support in previously untreated high-risk and older patients with CLL^32^. This combination was approved as a frontline treatment of CLL by the European Medicines Agency (EMA), based on the phase 3 clinical study^33^. In our CLL cohort, venetoclax + ibrutinib combination demonstrated a significantly elevated predicted synergy compared with random drug combinations (*p* < 0.0001; **Fig. 5E-F**), further supporting the ability of DANN to generalize across hematological malignancies and to recover clinically successful combination strategies.

We further investigated other than venetoclax-based combination regimens (**Suppl. Table 5**) by finding structurally similar compounds for four clinically approved drugs (cedazuridine, daunorubicin, glasdegib and ivosidenib). Among these, daunorubicin resulted in two compounds (CHEBI:94798 and NSC256439) with high enough structural similarity scores (Tanimoto>0.9) with compounds in the L1000 dataset. In the AML samples, both CHEBI:94798 + cytarabine and NSC256439 + cytarabine combinations showed significantly higher predicted synergy scores compared with random drug combinations (*p* < 0.0001; **Suppl. Fig. 11A–D**). In contrast, both of these combinations exhibited relatively low predicted synergy scores in the CLL samples, comparable to or even lower than those observed for many randomly selected combinations (**Suppl. Fig. 12A–D**). These treatments demonstrate disease-specific synergies, instead of broadly active combination effects.

Across both disease contexts, randomly paired drug combinations exhibited more broad and heterogeneous synergy distributions, often spanning both positive and negative values. In contrast, the clinically successful combinations showed a consistently more positive and concentrated synergy distribution, even though there were differences between the patients and synergy scores, as expected in such heterogeneous diseases (**Fig. 5, Suppl. Figs. 7-12**). The heterogeneity in combination responses was further reflected in the predicted errors and confidence scores of the combination responses between the patients, guiding a patient-specific tailoring of combinations that show not only high predicted synergy but also high prediction confidence. These results indicate that the DANN model effectively captures the patient-specific synergistic potential of clinically established drug combinations, rather than reflecting non-specific scoring biases that may lead to toxic combination responses.

Collectively, these findings demonstrate that the predicted synergy scores derived from the model and baseline RNA-seq data of patients before treatment align with known therapeutic efficacy, supporting its ability to identify both pharmaceutically and clinically meaningful drug combinations.

## Discussion

We developed a DANN framework for predicting drug combination synergy in individual leukemia patient samples, with the goal of improving generalization from preclinical cell line data to patient-derived samples. Our results demonstrate that domain-adversarial learning provides consistent improvements in predictive performance across multiple synergy metrics and disease contexts, particularly under limited data available from scarce patient samples. A central observation was that drug synergy distributions differ substantially between cell lines and patient samples. Across AML and CLL patients, patient-derived synergy scores exhibited broader dispersion and systematic shifts relative to cell line data. This discrepancy affects not only input feature distributions but also the statistical properties of the prediction target itself. Conventional machine learning models trained on cell line data implicitly assume stationarity of the outcome distributions, which leads to biased estimates when applied to patient contexts. By incorporating adversarial domain alignment during patient fine-tuning, DANN mitigates this mismatch and yields more robust cross-domain performance, hence advancing the development of predictive methods that are aligned with the limited sample sizes and other translational requirements of precision medicine.

In AML samples, we evaluated generalization across biological compartments by fine-tuning on BM samples and testing on PB samples. While all models showed improved performance as training sample size increased, as was expected, DANN consistently outperformed the non-adaptive baselines in the cross-compartment prediction, particularly in low-data settings. Importantly, this improvement was achieved without sacrificing within-domain performance on the training data, indicating that adversarial alignment did not compromise task-specific learning. Similar trends were observed across multiple synergy metrics, including HSA, Bliss and ZIP scores, demonstrating that the benefits were not specific to any single definition of drug interaction. These improvements were achieved by pre-training the model using combination data from only nine blood cancer cell lines, of which only a single cell line was originally derived from a patient with AML (**Suppl. Table 6**). The prediction results are therefore expected to improve in future with the availability of large-scale combination screens in cell lines and patient samples^34,35^. The DANN predictions showed an elevated synergy among venetoclax-based and other clinically approved combination regimens, supporting its ability to identify both pharmaceutically and clinically meaningful combinations.

We further demonstrated that these advantages generalize beyond AML to an independent CLL patient cohort, even though no CLL cell lines were used as part of model pre-training. Absolute predictive performance remained relatively moderate, reflecting the limited training data, intrinsic variability and limited dynamic range of *ex vivo* synergy measurements of CLL samples, which led to lower synergy score extremes than in the AML samples; yet, DANN consistently achieved higher accuracies and lower prediction errors than the non-adaptive baseline methods. These results indicate that pre-training on other blood cell line data can partly compensate for limited cell line and patient data, especially when combined with appropriate domain adaptation, which captures pharmacogenomic relationships that are transferable across leukemia types. Beyond predictive accuracy, we addressed prediction reliability through an error-guided selective prediction strategy. By training a *post-hoc* error model that estimates expected prediction error, we can prioritize high-confidence predictions and substantially improve the model’s effective performance in both AML and CLL cohorts. Notably, the error model generalized across diseases without any modification, suggesting that uncertainty patterns reflect shared properties of cross-domain synergy prediction, rather than disease-specific effects. This selective prediction framework provides a practical approach for balancing prediction coverage and accuracy, which may be particularly relevant in exploratory or hypothesis-generating settings in translational applications.

Toward translational applications, the proposed framework has the potential to be applied to patients with leukemia within a functional precision medicine setting. Following pre-training and patient-level fine-tuning, the model can be used to predict drug combination synergy scores for individual leukemia patients using bulk RNA-seq data of samples taken before treatment. Predictions can be made both for clinically established regimens and candidate combinations that have not yet been experimentally or clinically evaluated. In this context, higher predicted synergy scores that show high prediction confidence indicate combinations that are more likely to produce enhanced therapeutic effects in each patient, thereby supporting the prioritization of treatment options for further development. Consistent with this rationale, the model assigned high predicted synergy and confidence to clinically successful combinations, supporting its ability to capture therapeutically relevant interactions. Such predictions can complement *ex vivo* drug sensitivity testing by extending the search space of combinations beyond those that are experimentally feasible in scarce patient samples. In line with a recent report^36^, this framework leverages patient-specific adaptation together with preclinical knowledge to enable individualized prediction of combination responses across a wide range of combinations. Our results suggest that the model may serve as a practical tool to support personalized combination therapy optimization in AML and CLL, while further prospective validation will be required to establish its clinical utility.

Despite these promising results, several limitations should be acknowledged. First, patient sample sizes and the availability of AML and CLL cell line drug combination data remain modest, which limits the achievable performance and complexity of patient-specific adaptation. While domain-adversarial learning improves robustness under data scarcity, which is often the case in patient-based applications, it does not fully overcome the limitation imposed by small and heterogeneous patient datasets. However, the framework will benefit from increased availability of drug combination screening datasets in both cell lines and patient samples, which is expected to result in more accurate patient-specific predictions. Second, although we demonstrated generalization across two very distinct hematological malignancies, the applicability of the model to solid tumors, where microenvironmental and spatial effects may be more pronounced, remains to be established, once large-scale datasets of drug combination testing in patient tumors become available. Third, while DANN addresses domain shift at the representation level, it does not explicitly model patient-specific biological mechanisms underlying drug synergy. As a result, predictions should be interpreted as statistical estimates rather than mechanistic explanations. Integrating additional biological constraints or causal modeling may further improve model interpretability and robustness.

Finally, the error prediction module, while effective at ranking predictions by expected reliability, does not provide calibrated probabilistic guarantees, and its use introduces an explicit trade-off between prediction coverage and performance.

In summary, this work demonstrates that domain-adversarial learning provides a principled and effective strategy for improving cross-domain drug synergy prediction in leukemia patient samples. By explicitly accounting for systematic differences between cell line and patient responses and incorporating uncertainty-aware filtering, the proposed framework advances the methodological foundations of clinically-actionable drug combination response predictions. While further validation and extension are required, particularly in larger and more diverse patient cohorts, these results highlight the potential of domain-adaptive machine learning to bridge preclinical data and clinical domains in precision oncology.

## Materials and Methods

### Drug combination datasets

*In vitro* cell line drug combination data were obtained from DrugComb^37^, comprising a relatively limited number of blood cancer cell lines (*n* = 9, **Suppl. Table 6**) with experimentally measured drug combination responses (2,150 unique drug combinations tested). These cell line data originate from multiple combination screens published as separate studies and integrated in DrugComb. The cell line data were used here exclusively for model pre-training to learn generalizable drug combinations–response patterns.

AML *ex vivo* drug combination measurements were provided through the BeatAML project, and included samples derived from both bone marrow (BM) (*n* = 72 samples, 55 unique drug combinations) and peripheral blood (PB) (*n* = 74 samples, 55 unique drug combinations). The BM samples were randomly subsampled at varying sizes for patient-specific fine-tuning of the model, with the remaining BM samples used for internal testing. PB samples from independent AML patients were used as a separate held-out test set to evaluate cross-compartment generalization from BM to PB.

CLL drug combination data consisted of *ex vivo* drug response measurements from primary patient PB samples (*n* = 62 samples, 90 unique drug combinations). We implemented the same evaluation protocol applied across all compared prediction models, where an increasing number of patient samples were used for model adaptation to monitor predictive accuracy in the remaining CLL samples.

Drug synergy was quantified using multiple commonly used metrics, including Highest Single Agent (HSA), Bliss independence, and Zero-Interaction Potency (ZIP)^38^. Positive values indicate synergy, and negative values antagonism, while values close to zero correspond to additive combination effects. The analyses were conducted separately for each synergy metric to ensure consistency and robustness of performance assessment. We used HSA as the synergy score in the main figures, following the independent drug action concept^3^.

### CLL patient drug combination and transcriptomic assays

Small-molecule inhibitors (Selleck Chemicals) were dissolved in DMSO at 10 mM and arrayed into 384-well plates using echo acoustic (Labcyte Echo655 Dispenser, Beckman Coulter) and digital dispensing (Certus Flex, GYGER). Each drug pair was tested in a five-point five-fold serial dilution concentration series of the base drug (10 µM to 0.016 µM, except where otherwise specified) together with a combination drug at a single concentration (between 100 nM and 400 nM) or DMSO at a fixed 0.2% DMSO concentration. Cryo-preserved primary mononuclear cells were thawed as previously described^39^, and plated at a density of 20,000 cells per well in RPMI 1640 medium (Gibco) supplemented with 10% heat-inactivated human serum from male AB clotted whole blood (Sigma-Aldrich), 2% 200 mM L-glutamine (Gibco) and 1% 10,000 U/mL penicillin/10,000 µg/mL streptomycin (Gibco). Cells were exposed to drug combinations for 48 hours at 37°C in 5% CO_2_. Cell viability was assessed using the ATP-based CellTiter-Glo assay (Promega). Luminescence was measured on a BioTek Synergy LX multimode microplate reader (Agilent). Measured values were corrected for incubation effects^40^, and normalized to DMSO-treated control wells on each plate to obtain relative cell viability for each drug and dose combination in each patient sample.

Transcriptomic data of CLL patient samples were obtained from a previous study^39^. RNA from primary PB specimens was isolated using the RNeasy mini kit (Qiagen) and the quality assessed on an 2100 Bioanalyzer (Agilent) with RNA integrity scores above 8. Libraries for bulk sequencing were prepared according to the TruSeq RNA sample preparation v2 protocol (Illumina) and sequenced on Illumina HiSeq 2000 machines (paired-end, 2-3 samples multiplexed per lane). Reads were aligned to the GRCh37.1/hg19 genome using STAR (version 2.3.0) with default parameters. Gene-level counts were quantified using ht-seq-count with default mode union and the genes without valid annotation or gene-length information were removed. For each gene, counts were first normalized by gene length to obtain reads per kilobase (RPK), followed by library-size normalization to transcripts per million (TPM). TPM values were further transformed using log_2_(TPM + 1). Only patients with both transcriptomic profiles and matched drug-combination screening data were retained, resulting in 62 CLL patient samples for the downstream modelling.

### AML patient drug combination and transcriptomic assays

Small-molecule inhibitors were dissolved in DMSO and arrayed into 384-well plates. Drug combination experiments were performed using predefined pairwise inhibitor combinations, similar to previous study^41,42^, where each drug pair was tested in a seven-point fixed molar concentration series, matching the concentration range used for Beat AML single-agent assays (10 μM to 0.0137 μM, except where otherwise specified). Primary mononuclear cells were plated within 24 hours of sample collection at a density of 10,000 cells per well in RPMI 1640 medium supplemented with 10% fetal bovine serum, L-glutamine, penicillin/streptomycin, and β-mercaptoethanol. Cells were exposed to drug combinations for 72 hours at 37°C in 5% CO_2_. Cell viability was assessed using the MTS assay (CellTiter96 AQueous One, Promega), and absorbance was measured at 490 nm. Raw absorbance values were background-corrected and normalized to untreated control wells to obtain relative cell viability for each drug and dose combination in each patient sample.

Transcriptomic data of AML patient samples were obtained from the BeatAML cohort^42,43^. The primary AML specimens (BM and PB samples) were profiled by bulk RNA sequencing using the Agilent SureSelect strand-specific RNA library preparation workflow and Illumina HiSeq 2500 paired-end sequencing. Reads were aligned to the GRCh37 reference genome with Subjunc, and gene-level counts were quantified using featureCounts based on Ensembl release 75 annotations. Expression values were filtered and normalized using conditional quantile normalization to generate GC-content-corrected log_2_RPKM values. In the present study, we used the processed BeatAML transcriptomic data for modeling.

### Molecular feature representation

Each drug combination was represented using a multi-modal feature vector that integrates chemical structure information, drug-induced transcriptional signatures, and pathway-level activity features (from bulk RNA-sequencing). Drug-induced transcriptional signatures were derived from the LINCS L1000 data resource^44^, which quantifies gene expression changes following the chemical perturbations. For each compound, transcriptional responses were represented as z-score–normalized gene expression values relative to control conditions. For informative and compact drug representations, we performed gene selection based on variability across compounds and calculated the variance of each gene across all drug-induced expression profiles and selected the genes with variance greater than one. This resulted in a subset of 1,006 highly variable genes. The final drug signature for each compound was defined as a 1,006-dimensional vector consisting of the z-scored transcriptional responses in the given cell line. For each drug pair, the corresponding L1000 signatures of the two individual compounds were concatenated to form a combined transcriptional perturbation signature.

Chemical structure information of the small-molecule compounds was encoded using MACCS 166-bit fingerprints, as originally defined in the MACCS key set^45^. Molecular structures were represented as SMILES strings and processed using the rcdk package (version 3.6.0), which provides an interface to the Chemistry Development Kit (CDK). For each compound, SMILES strings were parsed into molecular objects using the parse.smiles function. MACCS fingerprints were then computed using the get.fingerprint function with type “maccs” and fp.mode “bit”, resulting in fixed-length binary vectors indicating the presence or absence of the predefined substructure patterns. The resulting fingerprints were truncated to the standard 166-bit representation. For each drug combination, the MACCS fingerprints of the two constituent drugs were concatenated to form a joint structural feature vector.

To capture pathway-level functional effects, single-sample Gene Set Enrichment Analysis (ssGSEA) was applied to baseline gene expression profiles of cell lines and patient samples^46^. To ensure consistency across the domain-specific datasets, we restricted sample-level bulk RNA-seq analyses to a common set of 7,831 genes, defined as the intersection of genes shared across the cell line dataset, the BeatAML patient cohort, and the CLL patient cohort (gene list provided in **Suppl. Table 7**). The cell line transcriptomic data were obtained from a previous study^47^. ssGSEA was performed with predefined gene sets (c2.cp.v2023.1.Hs.symbols.gmt, **Suppl. Table 8**), and the gene set enrichment scores were computed independently for each sample or cell line using the same gene sets for both cell line and patient samples, ensuring consistency across domains. This resulted in pathway activity profiles representing the relative baseline activation level of biological pathways.

### Domain-adversarial neural network (DANN)

The DANN architecture comprised three components: a shared feature extractor, a regression head for drug synergy prediction, and a domain classifier for distinguishing cell lines from patient-derived samples (**Fig. 1A**). The shared feature extractor was implemented as a feed-forward neural network that projects the concatenated multi-modal input features into a low-dimensional latent representation. The regression head maps this latent representation to continuous drug synergy scores. In parallel, the domain classifier was trained as a binary classifier to predict the data origin (cell line versus patient sample) from the same latent features. To enable adversarial learning, a gradient reversal layer was inserted between the feature extractor and the domain classifier. During model training, this mechanism encourages the feature extractor to learn representations that are predictive of drug synergy while minimizing domain-specific information, thereby promoting domain-invariant feature learning across experimental settings.

### Two-stage training strategy

Model training was conducted using a two-stage strategy to decouple the learning of general drug synergy patterns from patient-specific domain adaptation.

#### Stage I: Pre-training on cell line data

In the first stage, the model was trained exclusively on drug combination data from cancer cell lines. During this phase, only the shared feature extractor and the regression head were optimized, while the domain classifier was not used. The objective of this stage was to learn a general mapping between multi-modal molecular features and drug synergy outcomes under controlled experimental conditions. No patient-derived samples were included during pre-training. This design aims to ensure that the initial model captures broadly transferable drug response relationships without being influenced by patient-specific heterogeneity.

#### Stage II: Patient-specific domain adaptation

In the second stage, the pre-trained model was adapted to patient-derived data using a domain-adversarial learning framework. During this phase, patient drug combination measurements were used to supervise the regression head, while a domain classifier was introduced to distinguish between cell line and patient samples based on the shared latent representation. A gradient reversal mechanism was applied between the shared feature extractor and the domain classifier. As a result, the shared feature extractor was optimized jointly to (i) minimize prediction error on patient synergy measurements and (ii) reduce domain-specific information that allows discrimination between cell line and patient samples. This adversarial objective promotes the learning of representations that are predictive of drug synergy while being invariant to experimental domain differences.

#### Disease-specific adaptation

Domain adaptation and fine-tuning were performed separately for AML and CLL patient samples. This separation accounts for disease-specific biological characteristics, differences in data distributions and experimental conditions, and avoids conflating distinct synergy landscapes across diseases.

### Baseline comparison models

The DANN model was compared against several commonly used machine-learning baseline models for drug response prediction, including extreme gradient boosting (XGBoost)^48^, random forest (RF)^49^, k-nearest neighbors (KNN)^50^, and linear regression (LR)^50^. All baseline models were trained using the same multi-modal input features as DANN to ensure a fair comparison. Baseline models were trained directly on patient-derived data without domain-adversarial adaptation. For each task, identical training-test splits were applied across the DANN and the baseline models to enable direct comparison of their results.

### Model performance evaluation

Predictive performance was assessed by comparing predicted and measured drug synergy scores using Pearson correlation, Spearman correlation, and mean squared error (MSE). In AML patients, the model performance was evaluated in two settings: (i) on BM samples used during patient-specific fine-tuning and (ii) on held-out PB samples from independent patients, enabling assessment of cross-compartment generalization. In CLL patients, the performance was evaluated on held-out patient samples following the same evaluation protocol.

To assess robustness under limited patient data availability, models were trained using varying numbers of patient samples. For each training set size, random subsampling of patient samples was repeated five times. In each repetition, the DANN was re-initialized from scratch and trained independently, ensuring that no information was carried over across runs and preventing potential information leakage. Performance metrics were averaged across the five runs, and variability across the runs was quantified using standard errors.

### Ablation analysis of domain-adversarial learning

To evaluate the contribution of domain-adversarial learning, we conducted an ablation study in AML patient samples by comparing the full DANN model with a patient-only neural network lacking domain adaptation. The patient-only model shares the same network architecture as the task-specific predictor in DANN, but excludes the domain classifier and gradient reversal layer, and it was trained solely on BM patient samples. Both models were trained under identical experimental settings, including input features, training sample splits, and optimization procedures. Performance was evaluated on both BM samples used for training (within-domain) and PB samples held-out for testing (cross-compartment generalization). To ensure a fair comparison, we repeated the sampling of BM training subsets multiple times across varying sample sizes, and evaluated model performance using Pearson correlation, Spearman correlation, and MSE.

### Statistical evaluation of model performance via linear mixed-effects modeling

To evaluate the predictive performance of the DANN relative to baseline machine learning architectures, we utilized a linear mixed-effects modeling (LMM) framework. For each performance metric (Pearson correlation, Spearman correlation, and MSE), we specified a LMM wherein the metric value was modelled as a linear function of a ML method, sample size, and their interaction as fixed effects. Sample size was treated as a categorical factor to capture the non-linear trajectories of model performance. To control intra-iteration variability, the sampling iteration was incorporated as a random effect. The statistical significance of fixed effects was assessed using Type III Analysis of Variance (ANOVA), with Satterthwaite’s method for degrees of freedom approximation, as implemented in the lmerTest R package^51^. Subsequently, *post-hoc* pairwise comparisons of estimated marginal means were performed between DANN and baseline models using Tukey’s Honest Significant Difference (HSD) to control family-wise error rate. All statistical analyses were conducted in R (v4.4.1) using the lme4, emmeans, and broom.mixed libraries.

### Error prediction and uncertainty evaluation

To estimate prediction uncertainty and identify more reliable predictions, we trained an error prediction model to estimate the expected prediction error within DANN for individual patient samples. The error model was implemented as a random forest regressor and trained on patient-derived data using a three-fold cross-validation strategy to prevent information leakage between error estimation and performance evaluation. For each training split, patient samples were partitioned into a calibration subset and a disjoint evaluation subset. DANN predictions generated on the calibration subset were used to train the error model, ensuring that error estimates were learned without access to test set outcomes. Input features for the error model consisted of the same scaled multi-modal feature representation used by the DANN, augmented with the DANN-predicted synergy score and simple non-linear transformations of this prediction.

The target variable for error prediction was defined as the logarithm of the absolute prediction error, computed as the absolute difference between the predicted and measured synergy values followed by log transformation. This formulation stabilizes the error distribution and emphasizes relative rather than absolute deviations. During evaluation, the trained error model was applied to the test set to generate predicted error scores for each individual patient sample. The error scores were used to rank predictions by estimated uncertainty. Selective evaluation was then performed by retaining only a specified fraction of test samples with the lowest predicted error (the “kept ratio”), and recomputing performance metrics on the retained subset. Statistical significance of the performance differences across kept ratios was assessed using the Wilcoxon signed-rank test across repeated subsampling runs.

### Pathways analyses of combination responses

To investigate biological programs associated with drug combination responses, pathway-level analyses were performed by correlating pathway activity scores with observed drug synergy. Pathway activity scores were derived using ssGSEA as described above, based on baseline gene expression profiles of patient samples. For each drug combination, pathway activity scores were matched to the corresponding samples with measured synergy scores. Associations between pathway activity and drug combination synergy were assessed using Spearman correlation with two-sided significance testing, computed independently for each pathway. This analysis was performed separately for different sample types (BM and PB) to identify both shared and compartment-specific biological programs.

### Clinical validation of model predicted combinations

Clinically successful drug combinations in AML and CLL were identified from registered studies explicitly in ClinicalTrials.gov supporting FDA approvals for combination regimens (**Suppl. Table 5**), along with other clinically-used treatments (e.g., venetoclax + ibrutinib). FDA oncology-related approval notifications were examined through 2025 (inclusive), (https://www.fda.gov/drugs/drug-approvals-and-databases/resources-information-approved-drugs), and aggregated from two FDA resources (oncology/hematologic malignancy approval notifications and verified clinical benefit accelerated approvals). To ensure compatibility with the modeling framework and to allow comparison with random combinations, we verified that the selected clinically used drugs in combinations were present in both the L1000 drug perturbation dataset and the drug combination predictions for the AML and CLL patients.

Venetoclax combined with azacitidine represents a clinically established and widely used therapeutic strategy in AML, particularly in elderly patients or those unfit for intensive chemotherapy, where it serves as a standard-of-care regimen^52^. Therefore, evaluating whether the model can recapitulate the synergistic effects of this clinically used combination provides a critical test of its translational relevance. Since azacitidine is not included in the L1000 drug perturbation dataset, we identified structurally similar compounds within the L1000 library using MACCS fingerprint–based Tanimoto similarity. This approach enabled the selection of surrogate compounds that preserve key chemical features of azacitidine, hence allowing an indirect evaluation of the model’s ability to prioritize clinically relevant combination strategies. We defined Tanimoto > 0.9 as high enough structural similarity.

In addition to the surrogate compound analysis in AML patients using structurally similar compounds available in the L1000 dataset, we further extended the clinical validation framework to CLL patients by evaluating the clinically developed combination between ibrutinib and venetoclax, which has been recently approved as a frontline therapy based on phase 3 clinical trials^33^. To evaluate whether predicted synergy scores for clinically relevant combinations differ from background distributions, each predicted combination was compared with 10 randomly sampled drug pairs among those tested in the AML and CLL patient samples. Since many of the tested combinations are expected to lead to synergy, these random pairs do not provide a reference distribution of truly non-interacting drug pairs. Statistical significance was assessed using a two-sided Mann-Whitney U test.

## Data and code availability

- Codes, data and instructions for running the DANN model are available on GitHub: https://github.com/Jiezhu1995/Domain-adversarial-learning-predicts-clinically-actionable-drug-combination-synergy.
- All training and test datasets used for DANN and error prediction models are publicly available on Zenodo: https://zenodo.org/records/19749487.
- Unpublished drug combination screening data from leukemia patients have been made available to enable benchmarking and developing new prediction methods: https://github.com/Jiezhu1995/Domain-adversarial-learning-predicts-clinically-actionable-drug-combination-synergy/tree/main/data/patient.
- AML and CLL patient RNA-seq data were obtained from previous studies^39,42,43^.
- Cell line transcriptomic responses were obtained from the LINCS resource^44^.
- Cell line drug combination data were obtained from the DrugComb database^37^.
- Cell line bulk RNA-seq data were obtained from the Gene Expression Omnibus (GEO; accession GSE68379)^47^.
- All predictive models were implemented in Python using standard ML and DL libraries, including scikit-learn for classical models and processing, TensorFlow/Keras for neural network implementation, as well as for gradient boosting frameworks such as XGBoost. Neural networks were trained using stochastic gradient descent with Adam optimizer. Random seeds were fixed to ensure reproducibility across the runs.

## Acknowledgements

We thank all the patients who have participated in the studies, and authors of the publications who have made their data available. We thank Yingjia Chen for testing the DANN codes. CSC (IT Center for Science, Finland) for the computational resources. Funding support: T.A. Research Council of Finland (grants 344698, 345803, 367855, and 373493), under the frame of EP PerMed (CLL-CLUE and CLL-OUTCOME); the Cancer Foundation of Finland, the Norwegian Cancer Society (grant 273810), South-Eastern Norway Regional Health Authority (grants 2020026 and 2023105), and iCAN – Digital Precision Cancer Medicine Flagship (iCAN-MULTIDRUG). J.K. Sigrid Jusélius Foundation and Finnish Cultural Foundation. J.Z. China Scholarship Council (CSC) and Vilho, Yrjö and Kalle Väisälä Foundation. TZ. Clinical Research Priority Program ‘Next Generation Drug Response Profiling for Personalized Cancer Care’ of the University of Zurich; Swiss Cancer Research Foundation (grant KFS-4439-02-2018), Monique-Dornonville-de-la-Cour Foundation; EP PerMed (CLL-CLUE).

