## Supplementary Figures for "Domain-adversarial learning predicts clinically actionable drug combination synergy in leukemia patients using bulk transcriptomics data"

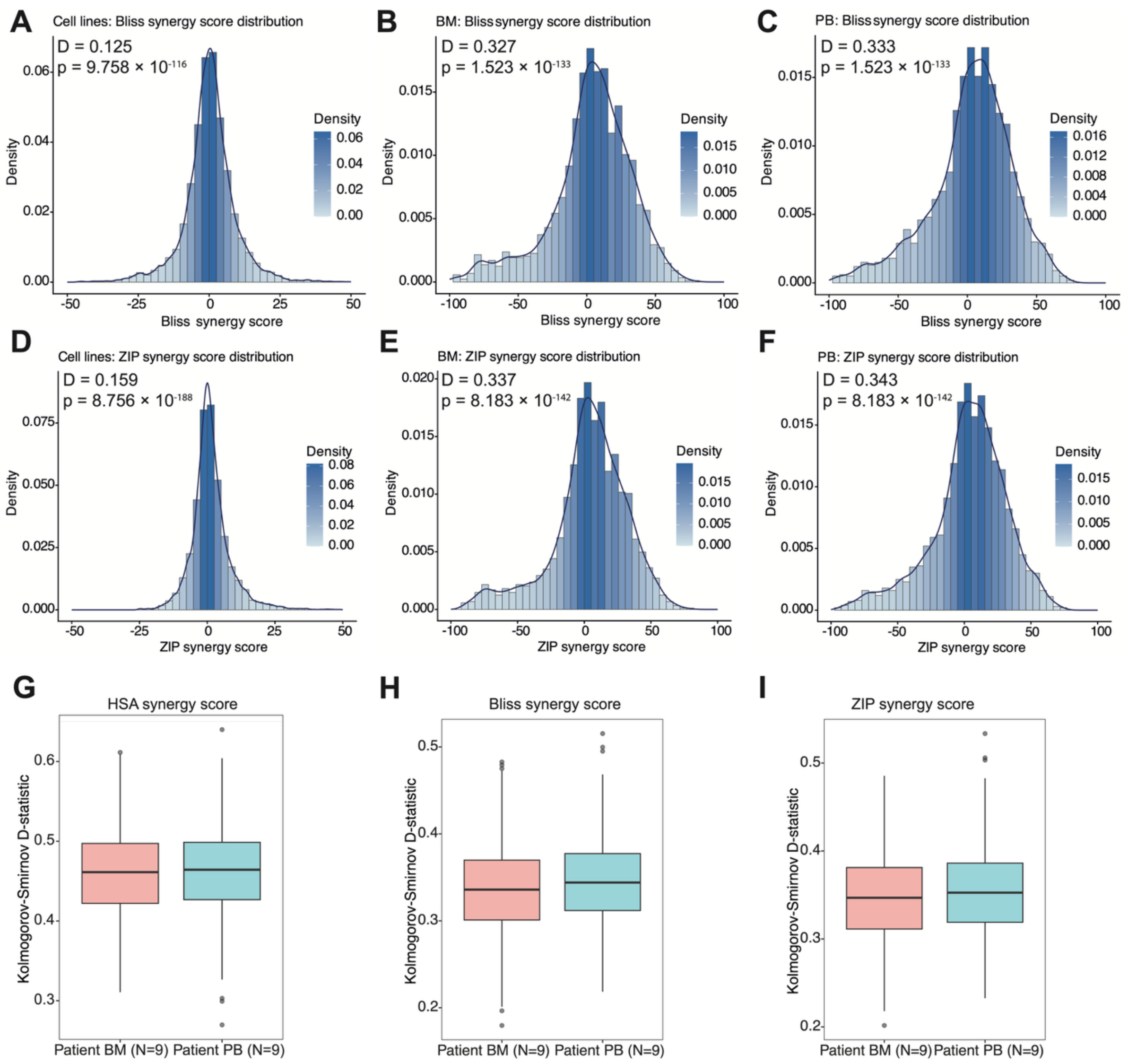


**Supplementary Figure 1. Distribution of Bliss and ZIP synergy scores across domains.** (**A**) Bliss synergy score distribution of 2,150 combinations tested in nine blood cancer cell lines, showing a relatively narrow range centered near zero. (**B**) Bliss synergy score distribution of 55 combinations in 72 AML BM patient samples, exhibiting a wider range of values. (**C**) Bliss synergy score distribution of 55 combinations in 74 AML PB patient samples, similarly showing increased dispersion compared with cell lines. (**D**) ZIP synergy score distribution of nine blood cancer cell lines, showing a relatively narrow range. (**E**) ZIP synergy score distribution in AML BM samples, with increased synergy variability. (**F**) ZIP synergy score distribution in AML PB samples, similarly spanning a broader range than cell lines. The distributional differences were assessed with the Kolmogorov–Smirnov test. (**G-I**) To address the sample size discrepancies between cell lines ($n$ = 9) and patient cohorts ($n$ = 72 or 74), a sub-sampling approach was run 1,000 times, where in each iteration nine patient samples were randomly selected for comparison against the cell line distribution. Across the three synergy metrics, consistently high D-statistics were observed, and all the patient subsamples showed statistically significant differences compared to cell line distribution ($p < 0.05$).

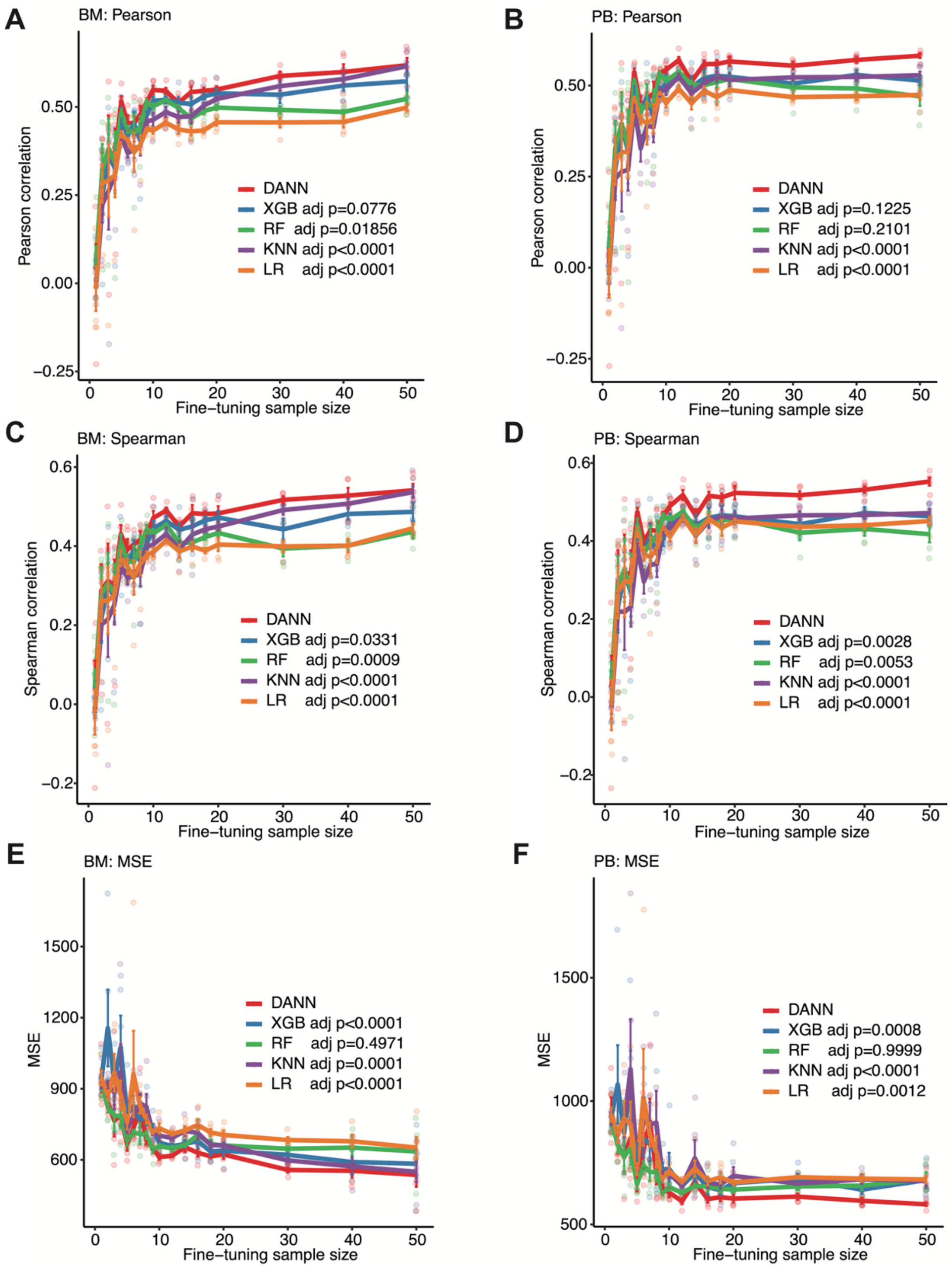


**Supplementary Figure 2. Predictive performance in AML samples using Bliss synergy scores.** Performance of DANN and baseline models on AML BM and PB samples, when the combination responses were evaluated using Bliss synergy scores. Panels correspond to Pearson correlation, Spearman correlation, and mean squared error as a function of training sample size, following the same evaluation protocol as in **Fig. 2**.

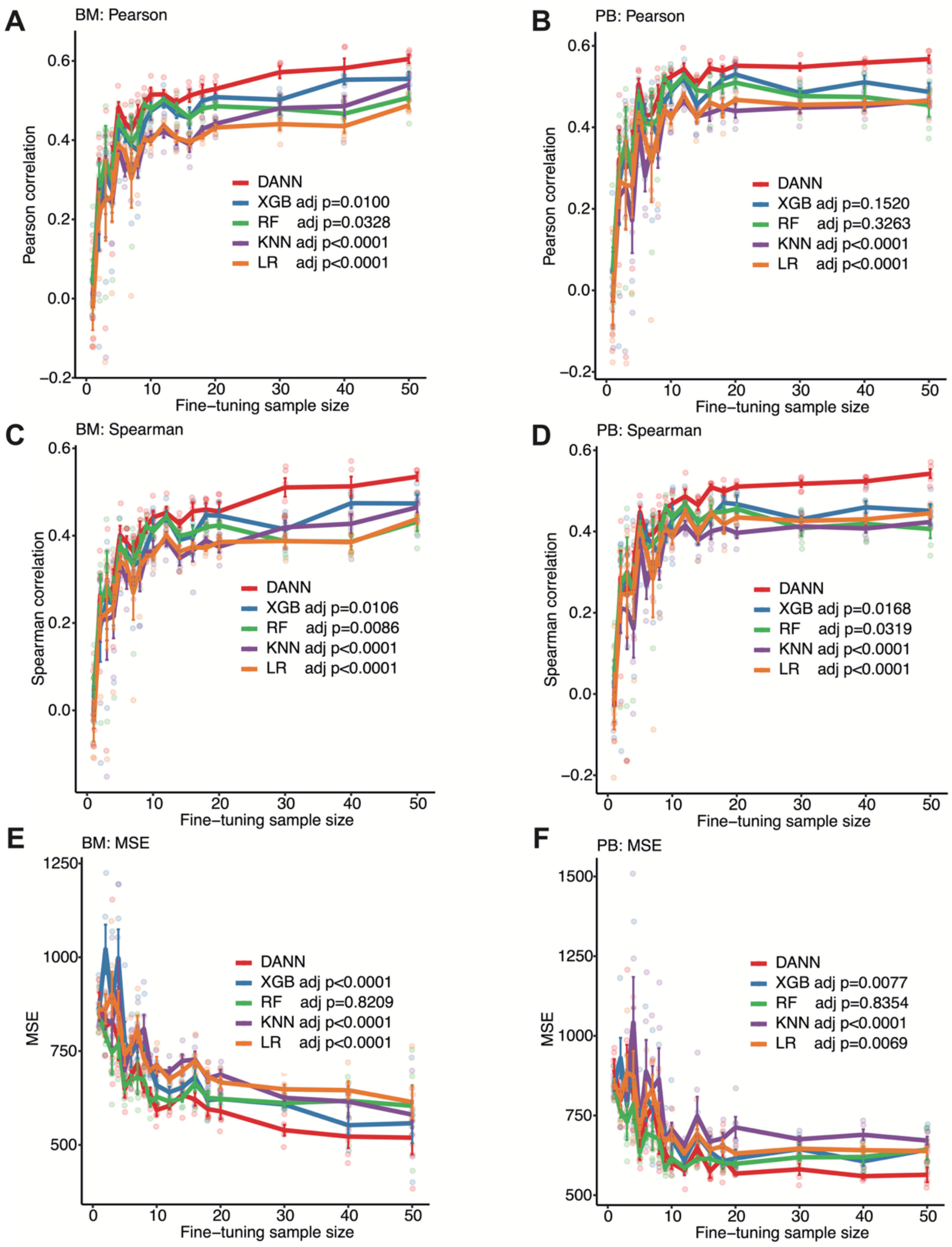


**Supplementary Figure 3. Predictive performance in AML samples using ZIP synergy scores.** Performance comparison of DANN and baseline models, when the combination responses were evaluated using ZIP synergy scores, reported for BM and PB samples across varying training sample sizes. Metrics include Pearson correlation, Spearman correlation, and mean squared error, analogous to the analyses shown for HSA and Bliss scores.

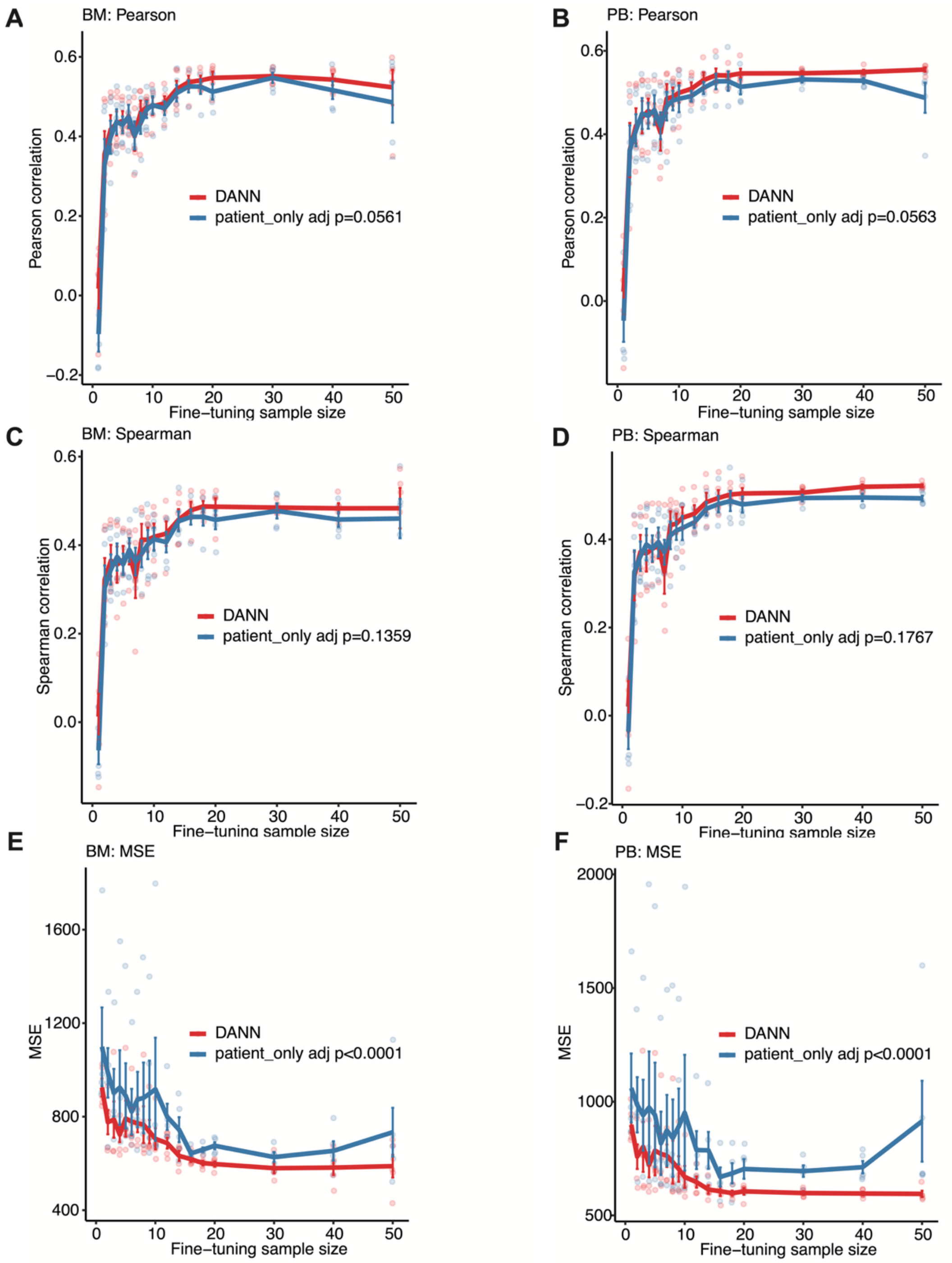


**Supplementary Figure 4. Ablation analysis of domain-adversarial learning in AML patient samples.** Predictive performance of the full DANN model compared with a patient-only neural network trained exclusively on BM samples without domain adaptation. Model performance is evaluated across increasing numbers of BM training samples and assessed on both the BM and PB datasets. (**A**, **B**) Spearman correlation on BM and PB samples. (**C**, **D**) Pearson correlation on BM and PB samples. (**E**, **F**) MSE on BM and PB samples. Points correspond to average performance across five repeated sampling runs, and error bars indicate standard errors. Significance was assessed via a robust LMM to account for inter-iteration variability. Pairwise contrasts between DANN and patient-only neural network model were adjusted for multiple testing using the Tukey's method (**Suppl. Table 3**).

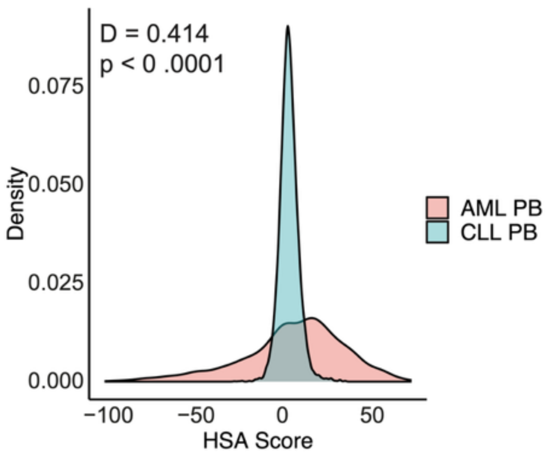


**Supplementary Figure 5. Distribution of HSA synergy scores in AML patient samples.** Density plots show the distribution of HSA synergy scores across the 74 AML PB samples (55 unique combinations) and 62 CLL PB samples (90 unique combinations). Significant distributional differences were observed between the disease contexts ($p < 0.0001$, Kolmogorov–Smirnov test), demonstrating a more compressed synergy landscape in the CLL cohort, likely due experimental design that used relatively low drug concentrations in the combination testing.

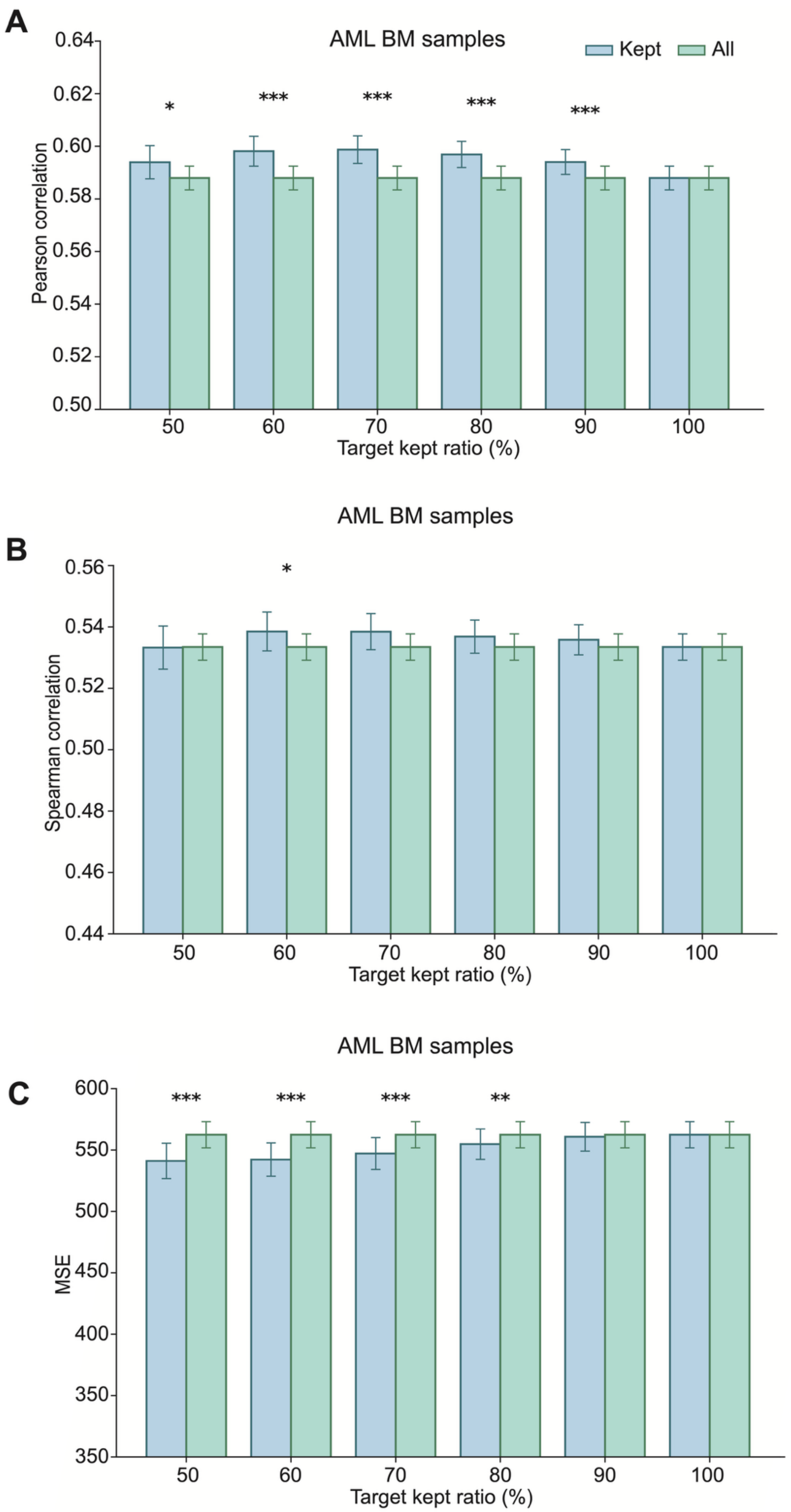
A
AML BM samples
Kept
All
Pearson correlation
0.64
0.62
0.60
0.58
0.56
0.54
0.52
0.50
*
***
***
***
***
50
60
70
80
90
100
Target kept ratio (%)
AML BM samples
B
Spearman correlation
0.56
0.54
0.52
0.50
0.48
0.46
0.44
*
50
60
70
80
90
100
Target kept ratio (%)
AML BM samples
C
MSE
600
550
500
450
400
350
350
***
***
***
**
50
60
70
80
90
100
Target kept ratio (%)

**Supplementary Figure 6. Error-guided selective prediction improves drug synergy prediction performance in AML bone marrow samples.** (**A**) Pearson correlation between predicted and observed synergy scores for retained ("kept") predictions compared with all predictions across different target kept ratios in 11 AML BM test samples. (**B**) Spearman correlation under the same selective filtering scheme. (**C**) MSE for kept versus raw predictions across target kept ratios. Predictions were ranked by estimated prediction error from the *post-hoc* error prediction model, and only the specified fraction of lowest-error predictions was retained for evaluation. Bars represent mean performance across repeated runs, with error bars indicating standard error. Statistical significance between kept and raw predictions at each kept ratio was assessed using the Wilcoxon signed-rank test (*$p < 0.05$, **$p < 0.01$, ***$p < 0.001$).

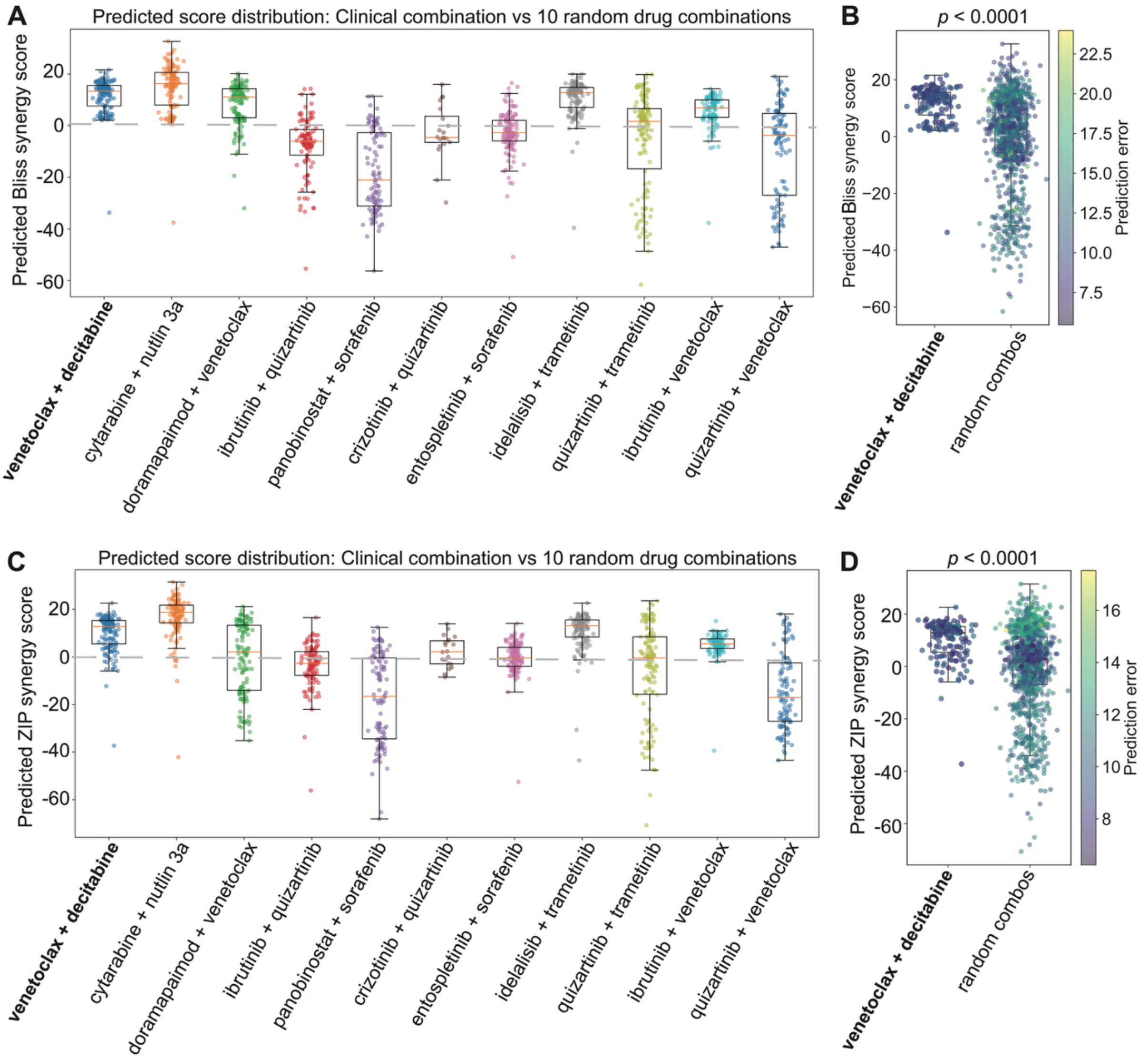


**Supplementary Figure 7. Predicted synergy scores for venetoclax + decitabine in the AML cohort.** (**A**, **C**) Distribution of predicted Bliss and ZIP synergy scores for the clinically relevant combination venetoclax + decitabine compared with 10 randomly sampled drug combinations in the AML cohort (*n* = 146). (**B**, **D**) Statistical comparison of predicted Bliss and ZIP synergy scores between venetoclax + decitabine and the aggregate of random drug combinations in the AML cohort, with significance assessed using a two-sided Mann–Whitney U test.

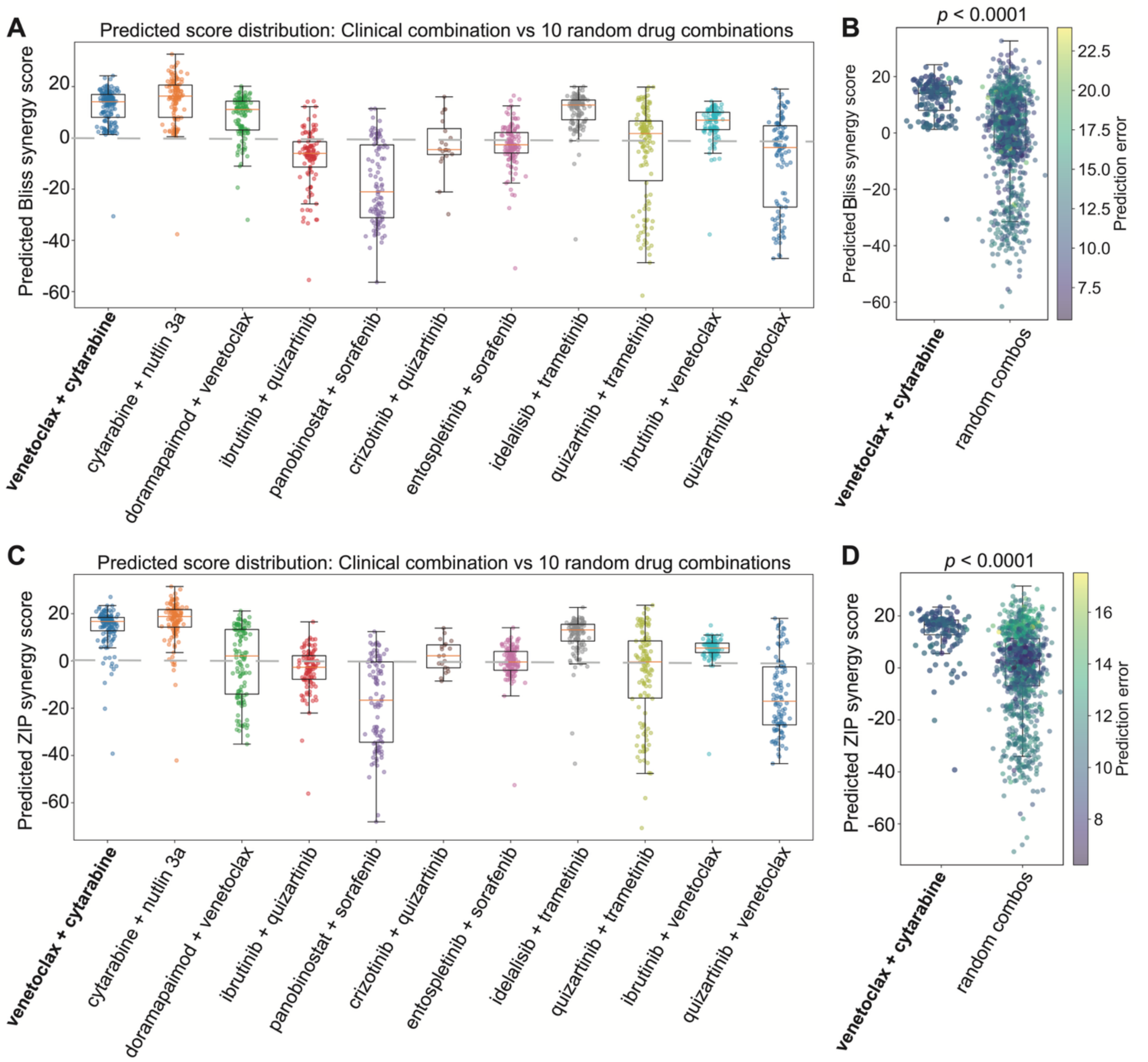


**Supplementary Figure 8. Predicted synergy scores for venetoclax + cytarabine in the AML cohort.** (**A**, **C**) Distribution of predicted Bliss and ZIP synergy scores for the clinically relevant combination venetoclax + cytarabine compared with 10 randomly sampled drug combinations in the AML cohort ($n$ = 146). (**B**, **D**) Statistical comparison of predicted Bliss and ZIP synergy scores between venetoclax + cytarabine and the aggregate of random drug combinations in the AML cohort, with significance assessed using a two-sided Mann–Whitney U test.

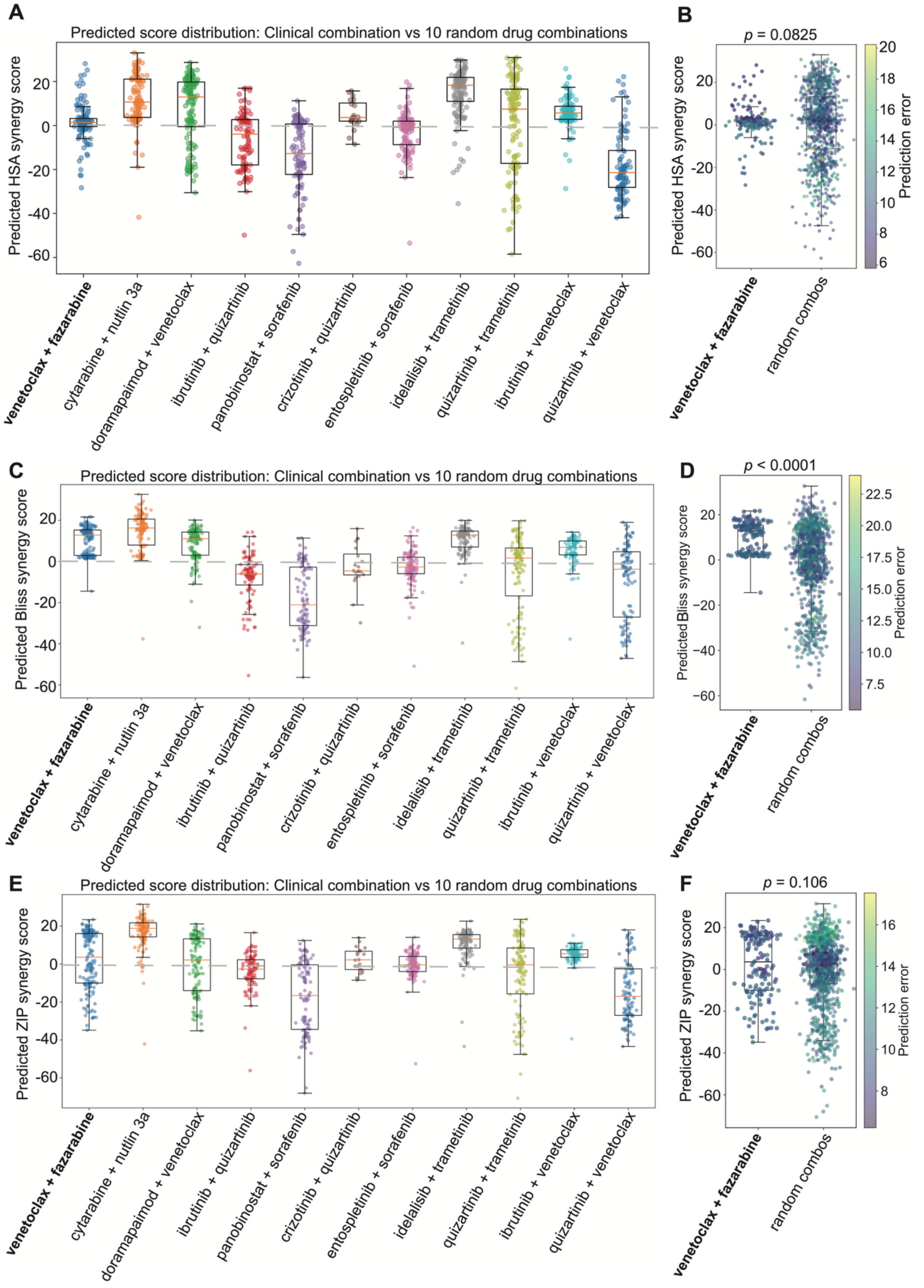


**Supplementary Figure 9. Predicted synergy scores for venetoclax + fazarabine in the AML cohort.** (**A**, **C**, **E**) Distribution of predicted HSA, Bliss and ZIP synergy scores for the clinically relevant combination venetoclax + fazarabine compared with 10 randomly sampled drug combinations in the

AML cohort ($n$ = 146). (**B**, **D**, **F**) Statistical comparison of predicted HSA, Bliss and ZIP synergy scores between venetoclax + fazarabine and the aggregate of random drug combinations in the AML cohort, with significance assessed using a two-sided Mann–Whitney U test.

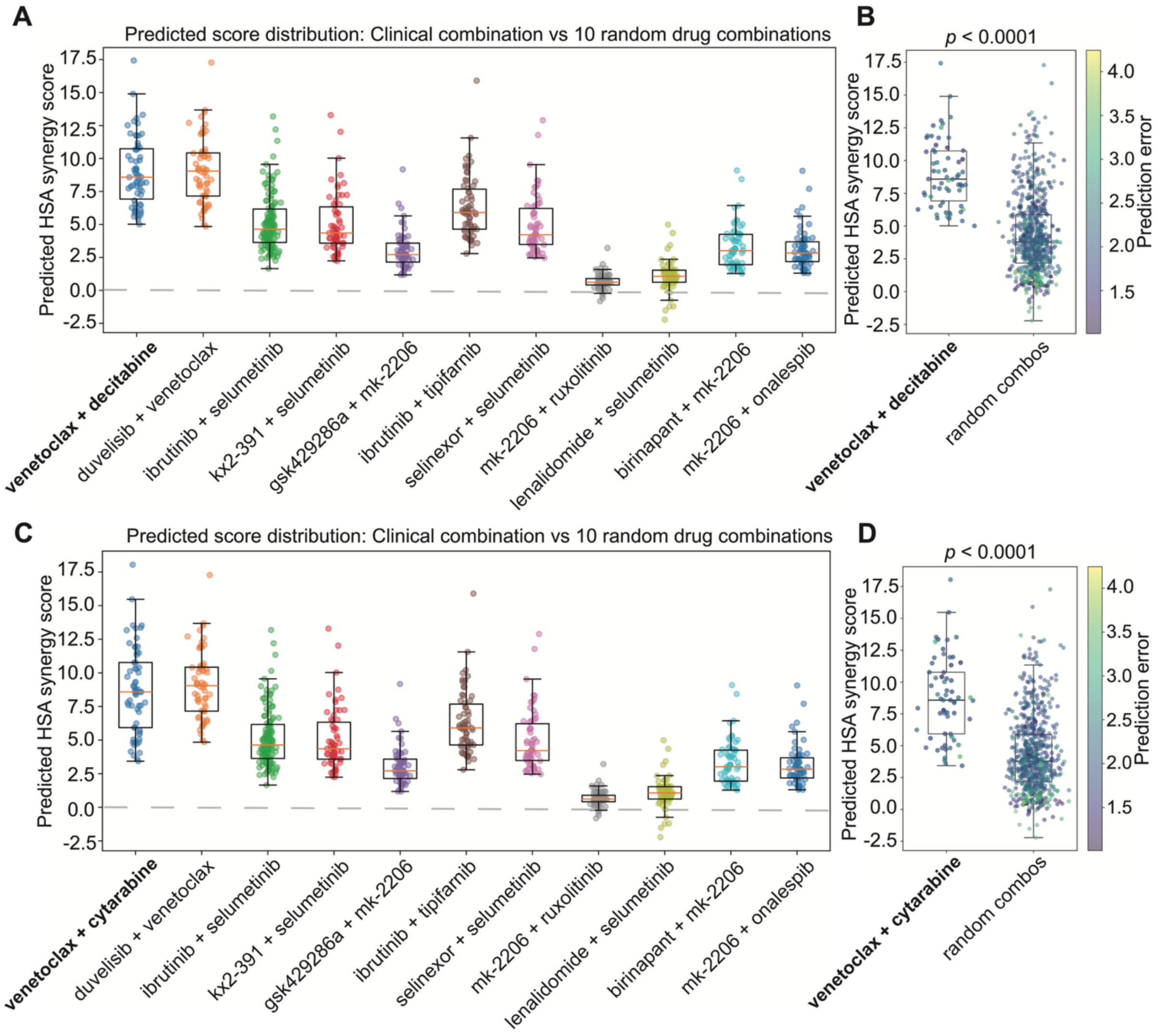


**Supplementary Figure 10. Predicted HSA synergy scores for clinically relevant combinations in the CLL cohort.** (**A**, **C**) Distribution of predicted HSA synergy scores for the clinically relevant combination: (**A**) venetoclax + decitabine and (**C**) venetoclax + cytarabine compared with 10 randomly sampled drug combinations in the CLL cohort ($n$ = 62). (**B**, **D**) Statistical comparison of predicted HSA synergy scores between each combination and the aggregate of random drug combinations in the CLL cohort, with significance assessed using a two-sided Mann–Whitney U test.

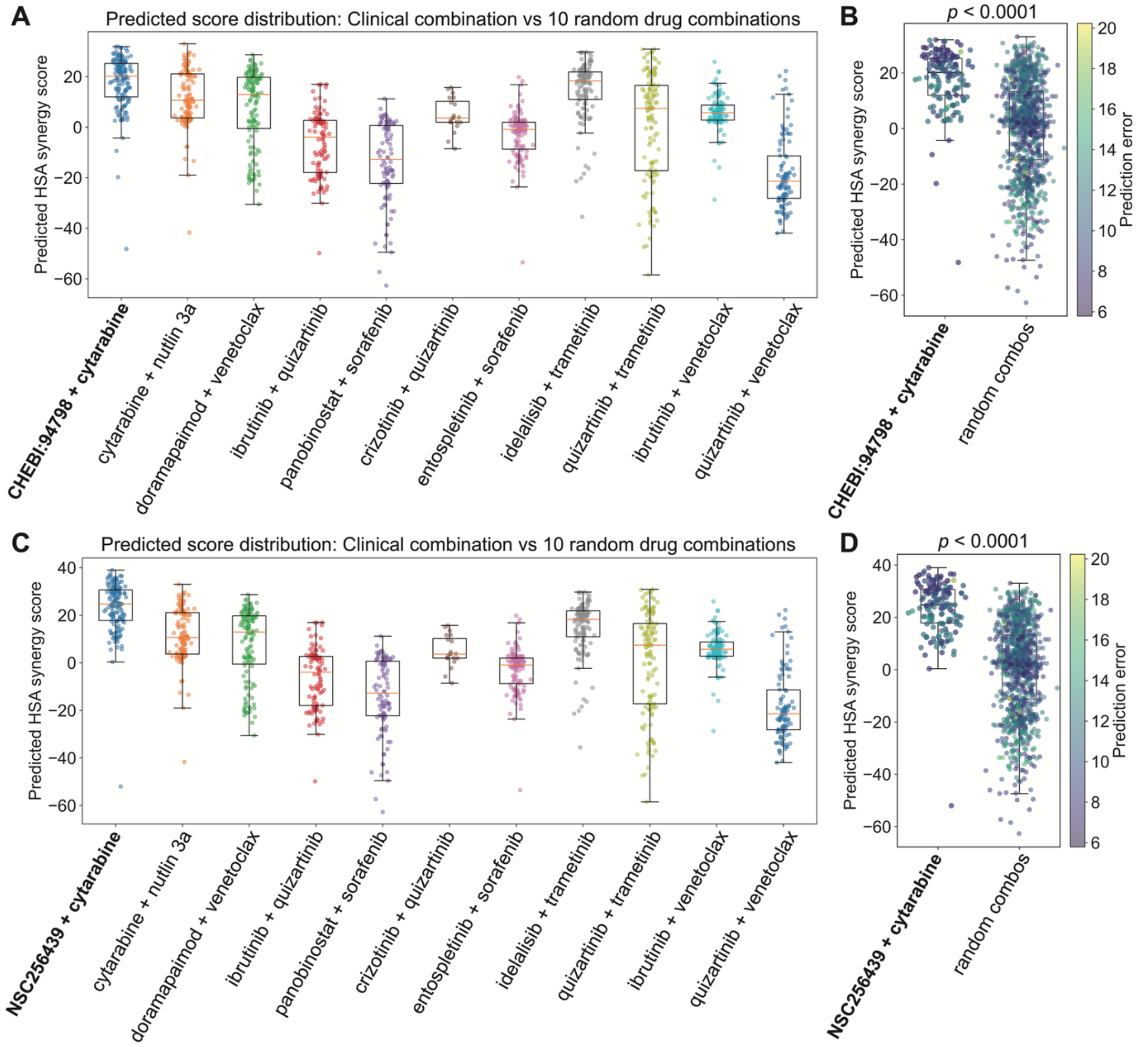


**Supplementary Figure 11. Predicted HSA synergy scores for daunorubicin + cytarabine structurally similar combinations in the AML cohort.** Distribution of predicted HSA synergy scores for the structurally similar daunorubicin-related combinations (bold): (**A**) CHEBI:94798 + cytarabine and (**C**) NSC256439 + cytarabine, compared with ten randomly sampled drug combinations tested within the AML cohort ($n$ = 146). Both structurally related combinations exhibit a shift toward higher predicted synergy values relative to the random background. (**B**, **D**) Statistical comparison of predicted HSA synergy scores between each structurally similar combination and the aggregate of random drug combinations in the AML cohort, with statistical significance assessed using a two-sided Mann–Whitney U test.

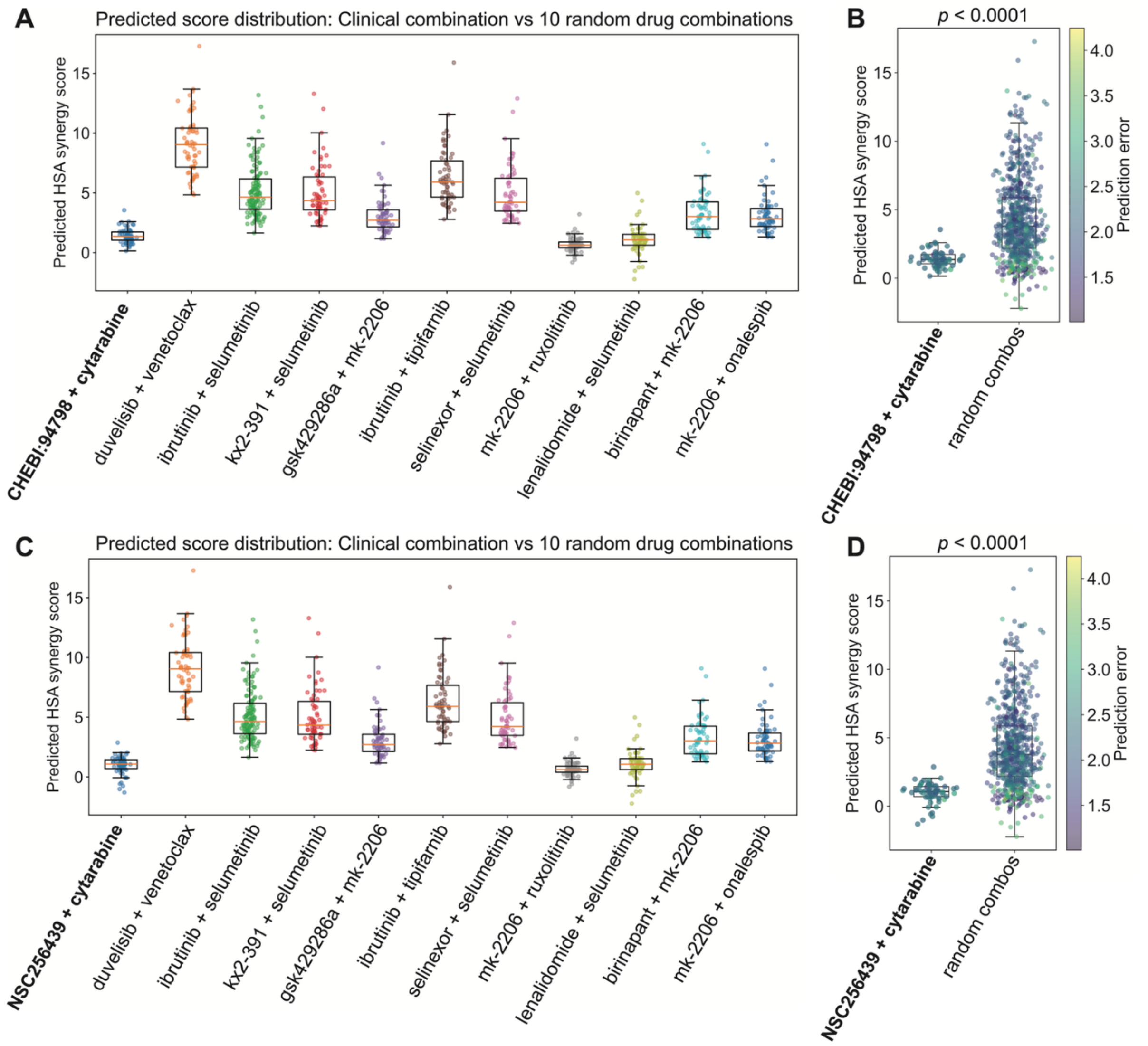


**Supplementary Figure 12. Predicted HSA synergy scores for daunorubicin + cytarabine structurally similar combinations in the CLL cohort.** Distribution of predicted HSA synergy scores for the structurally similar daunorubicin + cytarabine (bold): (**A**) CHEBI:94798 + cytarabine and (**C**) NSC256439 + cytarabine, compared with ten randomly sampled drug combinations tested within the CLL cohort. CHEBI:94798 and NSC256439 are structurally similar compounds to daunorubicin, a clinically established AML therapeutic commonly used in combination with cytarabine. In contrast to AML samples, both combinations exhibit relatively low predicted synergy values in the CLL cohort. (**B**, **D**) Statistical comparison of predicted HSA synergy scores between each structurally similar combination and the aggregate of random drug combinations in the CLL cohort, with statistical significance assessed using a two-sided Mann–Whitney U test.
